# Genomic and environmental controls on *Castellaniella* biogeography in an anthropogenically disturbed subsurface

**DOI:** 10.1101/2024.02.03.578758

**Authors:** Jennifer L. Goff, Elizabeth G. Szink, Konnor L. Durrence, Lauren M. Lui, Torben N. Nielsen, Jennifer V. Kuehl, Kristopher A. Hunt, John-Marc Chandonia, Jiawen Huang, Michael P. Thorgersen, Farris L. Poole, David A. Stahl, Romy Chakraborty, Adam M. Deutschbauer, Adam P. Arkin, Michael W. W. Adams

## Abstract

*Castellaniella* species have been isolated from a variety of mixed-waste environments including the nitrate and multiple metal contaminated subsurface at the Oak Ridge Reservation (ORR). Previous studies examining microbial community composition and nitrate removal at ORR during biostimulation efforts reported increased abundances of members of the *Castellaniella* genus concurrent to increased denitrification rates. Thus, we asked how genomic and abiotic factors control the *Castellaniella* biogeography at the site to understand how these factors may influence nitrate transformation in an anthropogenically impacted setting. ORR *Castellaniella* strains showed a higher degree of genetic diversification than those originating from non-ORR sites, which we attribute to the multitude of extreme stressors faced in the ORR subsurface. We report the isolation and characterization of several *Castellaniella* strains from the ORR subsurface. Five of these isolates match at 100% identity (at the 16S rRNA gene V4 region) to two *Castellaniella* amplicon sequence variants (ASVs), ASV1 and ASV2, that have persisted in the ORR subsurface for at least two decades. However, ASV2 has consistently higher relative abundance in samples taken from the site and was also the dominant blooming denitrifier population during a prior biostimulation effort. We found that the ASV2 representative strain has greater resistance to mixed metal stress than the ASV1 representative strains. We attribute this resistance, in part, to the large number of unique heavy metal resistance genes identified on a genomic island in the ASV2 representative genome. Additionally, we suggest that the relatively lower fitness of ASV1 may be connected to the loss of the nitrous oxide reductase (*nos*) operon (and associated nitrous oxide reductase activity) due to the insertion at this genomic locus of a mobile genetic element carrying copper resistance genes. This study demonstrates the value of integrating genomic, environmental, and phenotypic data to characterize the biogeography of key microorganisms in contaminated sites.

## INTRODUCTION

The nitrogen cycle has been profoundly impacted by agricultural and industrial activity, with excess nitrogen deposition contributing to the eutrophication of rivers, increased greenhouse gas emissions, and pollution of subsurface water sources (Ekemen Keskin, 2010, Kuypers *et al*., 2018). The Oak Ridge Reservation (ORR) in Oak Ridge, Tennessee is an extreme example of nitrate contamination, where concentrations in the subsurface reach up to 190 mM as a consequence of nuclear materials processing throughout the mid-20^th^ century (Thorgersen *et al*., 2019). The ORR Y-12 plant generated millions of liters of waste containing nitric acid and metals such as copper (Cu), cadmium (Cd), cobalt (Co), nickel (Ni), and uranium (U). Much of this liquid waste was deposited in an unlined surface disposal area (designated the S-3 ponds) between 1951 and 1983, resulting in extensive contamination of the subsurface (Brooks, 2001). Although attempts to remediate the site have been pursued, the subsurface environment surrounding the former S-3 ponds remains highly acidic and contains elevated levels of various toxic metals and nitrate (Brooks, 2001, Spain *et al*., 2007, Spain & Krumholz, 2011, Thorgersen *et al*., 2019). Despite the extreme conditions of the ORR subsurface, persistent microbial communities have been identified via metagenome surveillance and further characterized in culture-dependent studies (Spain *et al*., 2007, Green *et al*., 2010, Hemme *et al*., 2010). Community members include large proportions of nitrate reducing and denitrifying species (Green *et al*., 2010, Hemme *et al*., 2010, Spain & Krumholz, 2011). Thus, developing an understanding the abiotic and genomic controls on microbially driven nitrate reduction and nitrogen cycling at this site is important for predicting the fate of the contaminating nitrate.

A study examining the effects of electron donor injection on subsurface community structure and nitrate reduction activity at ORR found that biostimulation diminished overall microbial diversity, leading to an increase in Betaproteobacteria. Many of these Betaproteobacteria are capable of nitrate reduction, specifically members of the *Castellaniella* genus, suggesting that this genus may play a key role in nitrate remediation (Spain *et al*., 2007). *Castellaniella* strains have also been found in a variety of built and natural environments including forest soils (Foss *et al*., 1998), microbial fuel cells (Amanze *et al*., 2022), and sludges from wastewater treatment plants (Liu *et al*., 2008, Felföldi *et al*., 2010). These *Castellaniella* isolates are metabolically versatile and remarkably well-adapted for life in anthropogenically disturbed environments. For example, *Castellaniella* isolates have been shown to degrade cyclic monoterpenes under anoxic conditions (Petasch *et al*., 2014), as well as utilize taurine (Denger *et al*., 1997) and phenol (Ramezani & Zamir, 2022) as their sole electron donors under anoxic, nitrate-reducing conditions. Thus, this genus represents a relevant model for studying mechanisms of adaptive evolution and the influence of environmental stressors on diversification events.

The adaptive evolution of microbial genomes during periods of intense stress is proposed to be largely driven by horizontal gene transfer (HGT) (Hemme et al., 2016). HGT events generally involve the acquisition of mobile genetic elements (MGEs). MGEs can move between the genomes of two microorganisms or can move within the genome of a single organism. Specifically, comparative genomic analyses including pangenomic analysis allows for assessment of niche-specific adaptations (often horizontally acquired) that facilitate survival in contaminated environments (Gushgari-Doyle *et al*., 2022, Goff *et al*., 2022b). In this study, we present an analysis of *Castellaniella* diversity within the ORR subsurface within the broader context of other members of the *Castellaniella* genus. For this analysis we asked the following questions: *(i)* What is the geographic distribution of *Castellaniella* diversity in the ORR subsurface? *(ii)* Is the phylogenetic diversity of *Castellaniella* related to its broader habitat distribution? *(iii)* Does variation in phenotypes observed during laboratory simulations of relevant environmental stress explain the distribution and the differential abundance of distinct *Castellaniella* clades within the ORR subsurface? *(iv)* To what extent does horizontal gene transfer drive the ecological diversification of *Castellaniella*? To address these questions, we integrated environmental field, genomic, and phenotypic data to examine site-relevant traits of ORR *Castellaniella* isolates.

## MATERIALS AND METHODS

### Bacterial strains and growth conditions

Experimental work and genome sequencing was performed with six *Castellaniella* sp. strains isolated from groundwater samples collected at the Oak Ridge Reservation (ORR) in Oak Ridge, Tennessee, USA. *Castellaniella* sp. str. MT123 was previously isolated from groundwater samples collected from the contaminated well FW104 (lon. 35.97736048°, lat. –84.27356212°) adjacent to the former S-3 waste ponds at the Field Research Center in Oak Ridge, TN (Thorgersen et al., 2019). *Castellaniella* strains FW104-7C03, FW104-12G02, FW104-16D08, and FW104-7G2B were also isolated from this FW104 groundwater, and *Castellaniella* strain GW247-6E4 was isolated from groundwater taken from well GW247 (lon. 35.97729963°, lat. –84.27272624°) at ORR. Both wells are impacted by the contamination plume in the ORR subsurface (Smith *et al*., 2015). FW104-7C03, GW247-6E4, and FW104-7G2B were isolated on TSA plates under aerobic growth conditions at 30°C. FW104-12G02 was isolated on Eugon agar plates under aerobic growth conditions at 30°C. FW104-16D08 was isolated on LB agar plates under aerobic growth conditions at 30°C.

*Castellaniella* cultures were grown overnight by inoculating LB broth with five individual colonies grown on R2A plates. The overnight cultures were grown at 30°C while shaken at 200 rpm. For anaerobic growth experiments, 10 µL subsamples of the overnight cultures were inoculated into 400 μL of *Castellaniella* Experimental Medium (CEM) in individual wells of 100-well plates. For aerobic growth experiments, 5 µL subsamples of the overnight cultures were inoculated into 200 μL of *Castellaniella* Experimental Medium (CEM) in individual wells of 96-well plates. CEM contains, per liter, 0.6 g NaH_2_PO_4_, 20 mL of 1 M sodium lactate, 40 mL of a 25X salts solution, and 1 mL of a 1000X DL Vitamins stock and 10 mL of a 100X DL minerals stock as described previously in Widdel and Bak (1992). The 25X salts solution contains, per liter, 250 mg NaCl, 367 mg CaCl_2_ · 2H_2_O, 12.32 g MgSO_4_ · 7H_2_O, 2.5 g KCl, and 5 g NH_4_Cl. The pH of the medium was adjusted based on the desired conditions with HCl or NaOH. Different buffers were used based on the desired culture pH. For cultures which were grown at pH<7, we added 1 mL of 2 M sodium acetate buffer per liter. For cultures which were grown at pH≥7, a bicarbonate buffer was utilized. For the bicarbonate buffer, we added 2.5 g of NaHCO_3_ per liter. For anaerobic growth, cultures were amended with 10 mM NaNO_3_. For experiments monitoring the growth of *Castellaniella* isolates with metal exposure like that observed in well FW104, a metal mix representative of FW104 contamination (FW104 COMM) was included at 1X concentration in specified cultures. The final 1X FW104 COMM contained 215 μM Al(SO-_4_)_2_ · 12H_2_O, 80 µM C_4_H_6_O_6_U, 3000 µM MnCl_2_ · 4H_2_O, 20 µM NiCl_2_ · 6H_2_O, 2 µM CoCl_2_ · 6H_2_O, 1 µM CuCl_2_ · 2H_2_O, 10 µM Fe(SO_4_)(NH_4_)_2_(SO_4_) · 6H_2_O, and 1 µM Cd(CH_2_CO_2_)_2_ ·2H_2_O **(Table S1)**. The Al and U components were added from separate stock solutions, while the remaining metals were added from another 100X stock solution. This stock solution was stored in single-use aliquots at –20℃.

### Laboratory phenotyping of ORR isolates

Anaerobic growth experiments included the growth of *Castellaniella* isolates at various pH from 4 to 8, and their growth with or without the FW104 COMM. All anaerobic growth experiments were performed in 100-well plates in a Bioscreen incubating plate reader (Thermo Labsystems) placed in an anaerobic chamber under an 78% N_2_/20% CO_2_/2% H_2_ headspace. The Bioscreen monitored growth by optical density measurements at 600 nm (OD600) once every hour at 30°C. The Bioscreen was set to shake cultures continuously at low amplitude throughout the course of each growth experiment.

Aerobic growth experiments included the growth of *Castellaniella* isolates at pH 4 to 8. All aerobic growth experiments were performed in 96-well plates in a Cerillo Stratus plate reader placed in a shaking incubator set at 30°C. The plate reader read OD600 once every hour. The shaking incubator was set to shake continuously at 200 rpm throughout each growth experiment.

Nitrate reduction was assayed anaerobically using 10 mL of CEM containing 40 mM MES in Balch tubes with an 80% N_2_/20% CO_2_ headspace. Cultures were inoculated with 0.1 to 0.25 mL of overnight cultures and incubated as described above. Concentrations of nitrogen species were quantified as described previously (Goff *et al*., 2022b).

Analysis of growth curves was performed using the R (v4.2.3) package *gcplyr* (v1.5.2) as described in Blazanin (2023) with specific parameters described below. Prior to analysis of growth curve parameters, raw data were smoothed using the *moving-median* and *moving-average* smoothing algorithms. Additionally, OD600 values < 0.01 were excluded from further calculations to reduce the noise that arises at densities near 0. The per-capita growth rate (h^-1^) (*i.e*., the plain derivative divided by the population density) was calculated using the linear regression fitting functionality *window_width_n* of the *calc-deriv* algorithm with a window size of three data points.

### Curation and analysis of ORR amplicon sequence variant (ASV) data

Amplicon sequence variant data and associated geochemical data were retrieved from Goff *et al*. (2022b) and Ning *et al*. (2024). The Ning *et al*. (2024) dataset includes ASVs generated from a groundwater sampling survey performed in July and November of 2012 alongside nitrate concentrations, pH, and multiple heavy metal concentrations. These samples were collected from wells drilled at the site. We selected the subset of samples for our analysis collected from areas immediately adjacent to the former S-3 pond (referred to as “Area 3”, “Area 1”, and “Area 5”) **(Fig. S2).**

Sequencing was performed on a 10.0 µm fraction representing particulate-matter associated microorganisms and a 0.2 µm fraction representing planktonic microorganisms. The Goff *et al*. (2022b) dataset includes ASVs generated from a sediment core sampling survey performed in October of 2020 alongside porewater nitrate concentrations and sediment pH measurements (Putt *et al*., 2022). These samples all originated from within Area 3, immediately adjacent to the former S-3 ponds.

16S rRNA gene sequences were retrieved from the genomes of the six ORR *Castellaniella* isolates. These sequences were used to perform a BLASTn search (Altschul *et al*., 1990) against the ASVs from the two prior surveys. Additional *Castellaniella* ASVs were manually curated from the ASV matrix. Additionally, we identified samples from both prior surveys where *Castellaniella* ASVs were detected. For the 2020 sediment survey, relative abundances of ASVs were averaged across the two replicate samples. For the 2012 groundwater survey (Smith *et al*., 2015), relative abundances of ASVs were averaged between the two size fractions to consider the overall distribution of the ASVs at the site. To perform correlational analyses with geochemical parameters, we only considered the 0.2 µm fraction from the 2012 groundwater survey. Pearson correlation coefficients and p-values were calculated with R (v4.2.3) in RStudio (v2022.02.1 Build 461) using the function *rcorr* in the *Hmisc* package (v5.0-1).

### Genome sequencing and curation of publicly available genomes

High-molecular weight genomic DNA (gDNA) for MT123 was extracted using the Qiagen Genomic Tip 100/G kit according to the manufacturer’s protocol, except with the addition of 80uL of Qiagen lytic enzyme solution during the first digestion step with lysozyme. This same DNA was used as input to Illumina sequencing after needle shearing. The MT123 genome was sequenced by both Nanopore sequencing and Illumina sequencing as described previously (Goff *et al*., 2022a). Data processing for Nanopore reads and Illumina reads were performed as described in Goff *et al*. (2022a). Assembly was performed with the Nanopore and Illumina reads using Unicycler v0.4.8 as described in *Goff et al.* (2022a). The gDNA of isolates FW104-7C03, FW104-12G02, FW104-16D08, FW104-7G2B, and GW247-6E4 were extracted using a Qiagen DNeasy kit according to the manufacturer’s protocol for gram negative bacteria. Illumina Libraries were constructed with ∼250 ng of gDNA using an Illumina DNA Prep Tagmentation kits and Illumina DNA/RNA UD Indexes. Libraries were sequenced on an Illumina NovaSeq resulting in 2×150 paired end reads. The program Cutadapt v1.18 was used to remove adapter sequences (Martin, 2011), using the 3’ adapter sequence CTGTCTCTTATACACATCT. Sliding window quality filtering was performed with Trimmomatic v0.36 using parameters (-phred33 LEADING:3 TRAILING:3 SLIDINGWINDOW:5:20 MINLEN:50) (Bolger *et al*., 2014). All genomes were assembled de novo using SPAdes v3.15.3 with the following options (-k 21,33,55,77 –-careful) (Bankevich *et al*., 2012). Genome quality was validated with CheckM v1.0.18 using the lineage_wf pipeline with default parameters, maintaining only draft genomes with <10% contamination and >95% completeness (Parks *et al*., 2015). The above programs were run using the US Department of Energy Knowledgebase (KBase) (Arkin *et al*., 2018) using the KBase applications kb_cutadapt (v1.0.8), kb_trimmomatic (v1.2.13), kb_SPAdes (v1.3.3), and kb_Msuite (v1.4.0). All genomes were then annotated using the “Annotate Multiple Microbial Assemblies with RASTtk” (v1.073) KBase application using default parameters (Aziz *et al*., 2008, Overbeek *et al*., 2014, Brettin *et al*., 2015).

Eight high-quality draft and completed *Castellaniella* sp. genomes were found through the NCBI and Joint Genome Institute (JGI) Integrated Microbial Genomes & Microbiomes (IMG/M) databases. This was the total number of high-quality genomes that were available as of September 2022. Genome quality was determined using the BV-BRC Comprehensive Genome Analysis Service (Olson *et al*., 2023). Genomes annotated as “Good” quality by this tool were used for further analysis. Metadata including assembly length, isolation source, and sequencing information was also downloaded from IMG/M. The publicly available sequences were reannotated using the “Annotate Multiple Microbial Assemblies with RASTtk” (v1.073) application in KBase (Arkin *et al*., 2018) using default parameters (Aziz *et al*., 2008, Overbeek *et al*., 2014, Brettin *et al*., 2015).

### Phylogenetic analysis

To construct a phylogenetic tree that incorporates the ORR *Castellaniella* ASVs **(Table S2)** identified in the ORR environmental samples (16S rRNA gene V4 region, 253 bp), we extracted complete 16S sequences from the sequenced Castellaniella genomes. Note that the three metagenome-assembled genomes did not contain usable 16S rRNA gene sequences for this analysis. First, an alignment was constructed with the ASVs and full-length 16S rRNA gene sequences using MUSCLE (v3.8.425) in Geneious Prime with default settings. Alignments were trimmed to the 253 bp partial sequence. This trimmed alignment was used to generate a maximum likelihood tree with PhyML (v3.3.20180214) (Guindon *et al*., 2010) using an HYK85 nucleotide substitution model in Geneious Prime (v2022.0.2). Branch support was determined with 1000 bootstraps. Otherwise, default parameters were used. To determine the root of this tree, a second tree was constructed using the same method but including six outgroup 16S rRNA gene sequences from *Advenella mimigardefordensis* DPN7, *Alcaligenes faecalis* subsp. faecalis NBRC 13111, *Pusilimonas noertemannii* BS8, *Bordetella pertussis* Tohoma I, *Achromobacter oxylosoxidans* NBRC 15126, and *Bordetella petrii* DSM 12804. The original tree was re-rooted based on the results of this second tree containing the outgroup sequences. All phylogenetic trees in this study were visualized using iToL (Letunic & Bork, 2021).

A multi-locus tree was constructed in KBase with the 14 *Castellaniella* genomes using the Insert Set of Genomes Into SpeciesTree application (v2.2.0) (Price *et al*., 2010). The marker sequences for building the concatenated alignment are given in **Table S3**. Average Nucleotide Identity (ANI) was also calculated for these genomes using the Compute ANI with the FastANI (Jain *et al*., 2018) application in KBase.

### Pangenome Analysis

A pangenome comparing eight publicly available *Castellaniella* sp. genomes with six additional genomes presented herein (for a total of 14 genomes) was computed in KBase using the “Compute Pangenome” (v0.0.7) application with default methods. Genes were categorized as part of core (present in 100% of genomes), shell (present in 16-99% of genomes) and cloud (present in less than 1-15% of genomes). The openness of the pangenome was calculated using power-law regression (G = cN^γ^) as previously described (Liao *et al*., 2021) where G represents the pangenome size, c represents the core genome size, N is the number of genomes included in the pangenome analysis, and γ is the openness coefficient.

### Functional Annotations

The gene families from each genome were functionally characterized using the COG functional category with eggNOG-mapper (v2.1.9) under default parameters (Huerta-Cepas *et al*., 2019). The genomes were analyzed for heavy metal homeostasis genes (HMHGs) using Geneious Prime (v2022.0.2) to perform a BLASTp search against the BacMet Antibacterial Biocide and Metal Resistance Genes Predicted Database (v2) (Pal *et al*., 2013).

Curation of the results followed: (1) results with an e-value > 1E-10 were discarded; (2) results with a query coverage <70% were discarded; (3) results with a pairwise identity <25% were discarded; and (4) results with 25%< pairwise identity < 80% were manually assessed for likelihood of a positive hit based on sequence length, similarity of the protein annotation, and a BLASTP (Altschul *et al*., 1990) search against the UniProtKB/SwissProt database (Consortium, 2018). Manual review of annotations identified genome features associated with the denitrification process. Denitrifying genes were identified manually in accordance with previous reports (Tiedje, 1988, Kuypers *et al*., 2018).

### Chromosomally Integrated Element Analysis

The web-based software, IslandViewer 4 (Bertelli *et al*., 2017) was used to broadly identify genomic islands (integrating two different methods: SIGI-HMM (Waack *et al*., 2006) and IslandPath-DIMOB (Hsiao *et al*., 2003)). Further classification of IslandViewer4 predictions was conducted using PHage Search Tool Enhanced Release (PHASTER) (Zhou *et al*., 2011, Arndt *et al*., 2016) and ICEFinder (Liu *et al*., 2018) to identify prophage regions and integrative and conjugative elements, respectively.

### Castellaniella Global Geography

*Castellaniella* 16S rRNA gene sequences were retrieved from the SILVA database (v.138.1) in August 2023. Associated metadata were used to determine the environmental classification of and anthropogenic impact on the environmental sites of origin of these sequences. Our environmental classification scheme was based on the JGI GOLD ecosystem ontology (Mukherjee *et al*., 2020). For mapping, we extracted coordinates and/or location information from the metadata.

### Other Data Visualization

Heatmaps were visualized using the *pheatmap* (v1.0.12) package (Kolde & Kolde, 2015) in R (v4.2.3) in RStudio (v2022.02.1 Build 461). Correlelograms were visualized using the *corrplot* (v0.92) R package. Other plots were generated using the *ggplot2* (v3.4.2) R package.

## RESULTS AND DISCUSSION

### *Castellaniella* diversity, distribution, and history in the ORR subsurface

Using 16S rRNA (V4 region) gene amplicon sequence variants (ASVs) obtained in prior community analyses (Smith *et al*., 2015, Goff *et al*., 2022b, Ning *et al*., 2024), we examined the geographic range of *Castellaniella* in the region of the ORR subsurface surrounding the former S-3 ponds **(Fig S1).** We detected 12 *Castellaniella* ASVs across these samples **(Table S2, Fig. 1A, B, Fig. S2)**. These ASVs represent the V4 region of the 16S rRNA gene. Of these 12 ASVs, three were found to exceed a relative abundance of 0.5% in one or more samples: ASV1, ASV2, and ASV12. These three ASVs were present to varying degrees in the earlier community surveys, with ASV2 reaching the highest average relative abundance at 4.3% of the well FW104 groundwater ASVs. To link genotypic and phenotypic analyses with field observations, we isolated six *Castellaniella* strains from the contaminated ORR groundwater: FW104-16D08, FW104-12G02, FW104-7G2B, FW104-7C03, GW247-6E4, and MT123 **(Table S4)**. Strain GW247-6E4 originated from the contaminated well GW247 while the remaining strains all originated from the contaminated well FW104. FW104 is the location of the *Castellaniella* bloom described above that was observed in the 2012 survey (Smith *et al*., 2015). The strain MT123 V4 region matches ASV2 at 100% nucleotide identity. The V4 region of strains FW104-16D08, FW104-12G02, FW104-7G2B, and FW104-7C03 match with 100% nucleotide identity to ASV1. The GW247-6E4 V4 region did not match at 100% identity to any of the 12 *Castellaniella* ASVs **(Fig. 1A, Fig. S2)**. While ASV12 is generally found at higher abundance than ASV1 **(Fig. 1A),** unfortunately we were unable to recover a cultured representative of this population.

**Figure 1.**
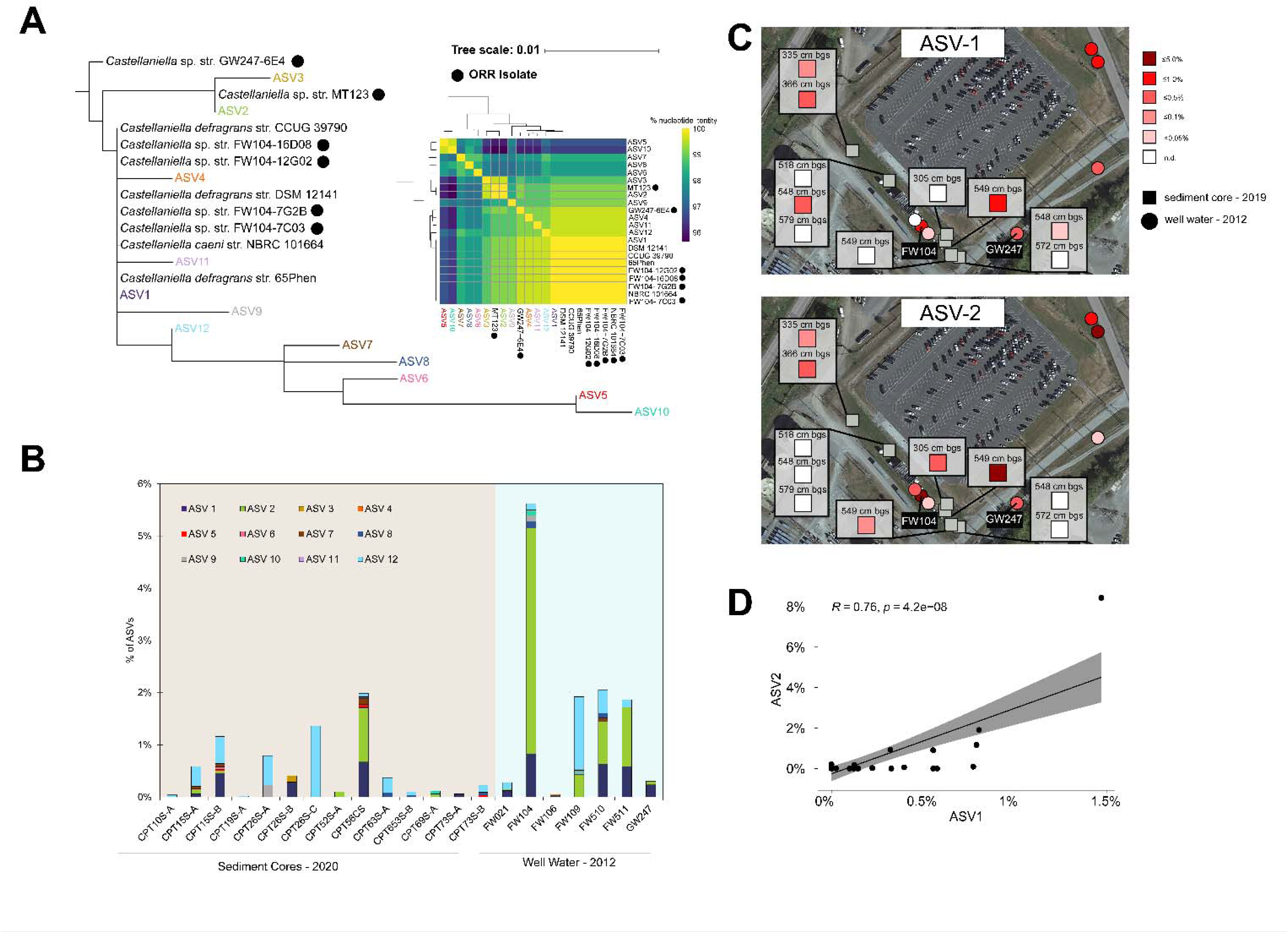
Ecology of ORR *Castellaniella* populations. (**A**) Maximum-likelihood tree of *Castellaniella* 16S rRNA gene V4 regions, incorporating the ORR ASVs (left) and a matrix showing the percentage nucleotide identity for the same set of sequences (right). ORR isolates are indicated with a black circle. The associated alignment is shown in **Fig. S2. (B).** Relative abundances of *Castellaniella* ASVs around the former S-3 ponds. Only samples with detectable *Castellaniella* ASVs are shown. **(C)** Spatial distributions in the ORR subsurface of two dominant ASVs with cultured representatives. Map data provided by Google Maps ©2023 Airbus, Maxar Technologies, U.S. Geological Surveym USDA/FPAC/GEO. **(D)** Correlation of ASV1 and ASV2 relative abundances across samples where *Castellaniella* are detectable. The trend line is shown in black, and the grey area reflects the uncertainty.

We next mapped the abundance patterns of the two ASVs with representative isolates (ASV1 and ASV2) **(Fig. 1C)**. As described above, there was a focal hotspot of both ASV1 and ASV2 in the FW104 groundwater in a 2012 groundwater survey (Smith *et al*., 2015). These two ASVs were also observed at higher abundance in a nearby sediment core during a later 2020 survey (Goff *et al*., 2022b) **(Fig. 1C)**. We found a positive correlation (*Pearson correlation coefficient = 0.75, p = 4.2E-6*) between the relative abundances of these two ASVs across all samples, suggesting that these two closely related clades stably coexist in the ORR subsurface (**Fig. 1D)**. Strong positive correlations were also observed between planktonic ASV1 and ASV2 abundances and Fe concentrations in the groundwater **(Fig. S3)**. Recently, we found that mixed metal exposure may significantly disrupt intracellular iron homeostasis, potentially increasing Fe demand in organisms at sites like the ORR (Goff *et al*., 2023). We observed negligible correlations between [Cu], [Cd], [U], [nitrate], and pH and the relative planktonic abundances of the two ASVs in the groundwater.

Biostimulation efforts conducted around the former S-3 ponds in the early 2000s induced *Castellaniella* blooms and enhanced denitrification rates (Spain *et al*., 2007). Spain *et al*. (2007) reported on a 16S rRNA gene clone library from an ORR sediment core collected after biostimulation with ethanol and bicarbonate. In this core, two *Castellaniella* clones dominated: “Operational Taxonomic Unit (OTU)34” (70% relative abundance) and “OTU35” (11% relative abundance). Comparisons to the V4 region of these clones revealed that ASV2 has 100% sequence identity to the V4 region of OTU34, except for a single nucleotide ambiguity in the OTU34 sequence **(Fig. S4)**. Additionally, the ASV2-representative MT123 16S rRNA gene sequence has 100% nucleotide identity to the full-length OTU34 sequence except for the single base pair ambiguity **(Fig. S5).** Thus, MT123 is a representative of a *Castellaniella* lineage that has persisted for at least two decades across multiple locations at this site, undergoing periodic blooms in response to modulation of its local environmental chemistry. We predicted that genomic features and laboratory phenotypes of these representative strains could be used to informs the controls on the distribution of these two dominant *Castellaniella* ASVs at the site.

### General features of *Castellaniella* genomes

We sequenced the genomes of the six ORR *Castellaniella* isolates. The MT123 genome was completed with long-read sequencing and the remaining are high-quality drafts (<60 contigs, N50 > 237,000). For additional context, we downloaded eight high-quality *Castellaniella* genomes and associated metadata from IMG and NCBI. These genomes originated from sediment, wastewater treatment bioreactors, and activated sludge **(Table S4).** Six of these genomes originated from sludges or bioreactors of wastewater treatment plants where physiological stressors such as high concentrations of antibiotics and nitrates are commonly found (Kümmerer *et al*., 2000, Fan *et al*., 2020). One of these genomes (FW021bin21) was a previously published metagenome-assembled genome (MAG) that originated from well FW021 which is adjacent to the former S-3 ponds at the ORR (Tian *et al*., 2020). The FW021bin21 MAG lacks a 16S sequence, and so we are unable to compare it to our database of ORR ASVs. No major differences in general genomic features (rRNA, tRNA, G+C content, and genome size) were observed when comparing ORR *Castellaniella* genomes to non-ORR *Castellaniella* genomes **(Fig. S6)**.

### A refined *Castellaniella* phylogeny using Average Nucleotide Identity (ANI)

ANI analysis for all *Castellaniella* genome pairs revealed six distinct clades with ANI values greater than 95%. We consider each of these clades to represent different *Castellaniella* species (Goris *et al*., 2007). This phylogeny is further supported by a multilocus species tree with all fourteen genomes **(Fig. 2A, B)**. While we had hypothesized that the genomes derived from ORR would cluster with each other given their shared origin, this was not the case. Based on the multilocus species tree and matrix of ANI values, only one clade contained multiple ORR genomes: ASV1 representatives FW104-16D08, FW104-12G02, FW104-7G2B, and FW104-7C03, all originating from FW104. The ASV2-representative genome, MT123, forms a clade with DR1149, originating from an anaerobic digester. Two other ORR genomes, FW021bin21 and GW247-6E4, formed singleton clades **(Fig. 2B).** Thus, despite originating in close physical proximity, the ORR *Castellaniella* genomes form four distinct clades, including three that likely represent novel *Castellaniella* species.

**Figure 2.**
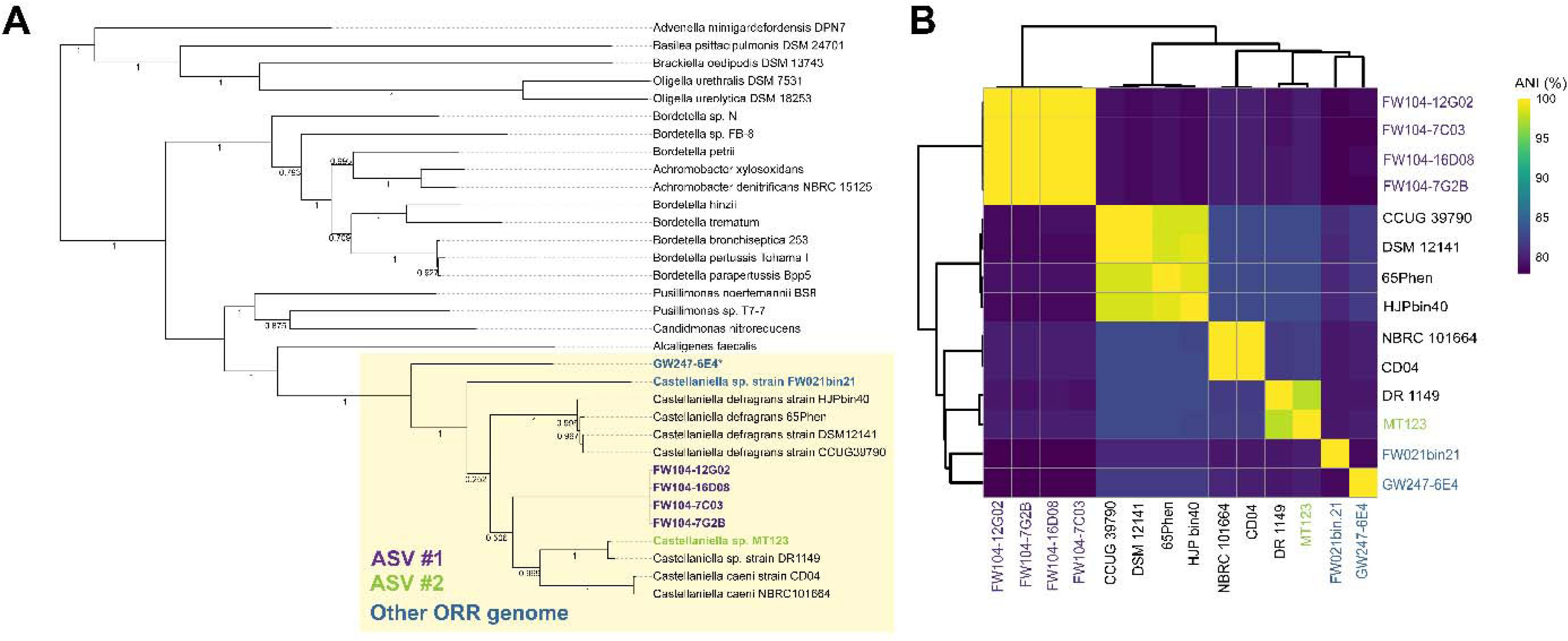
*Castellaniella* phylogeny. (**A**) Multi-locus phylogenomic tree of *Castellaniella* and close relatives using a concatenated alignment of 49 single-copy marker genes **(Table S3).** Bootstrap values are shown on the tree. **(B)** Average nucleotide identity (ANI) matrix of *Castellaniella* genomes clustered using a Euclidian distance metric.

### Pangenome structure

A *Castellaniella* pangenome was computed using the 14 genomes to further characterize the genetic diversity of this genus. The resulting pangenome contained 9,326 unique orthologous gene clusters (orthologs) that were categorized based on the percentage of genomes containing the given ortholog. The orthologs were categorized as core (present in 100% of genomes), shell (present in 16-99% of genomes) and cloud (present in less than 1-15% of genomes). The *Castellaniella* pangenome (n=14) contained 1,228 core gene clusters (13% of the pangenome), 3,291 shell gene clusters (35%) and 4,807 cloud gene clusters (51%) **(Fig. 3A)**.

**Figure 3.**
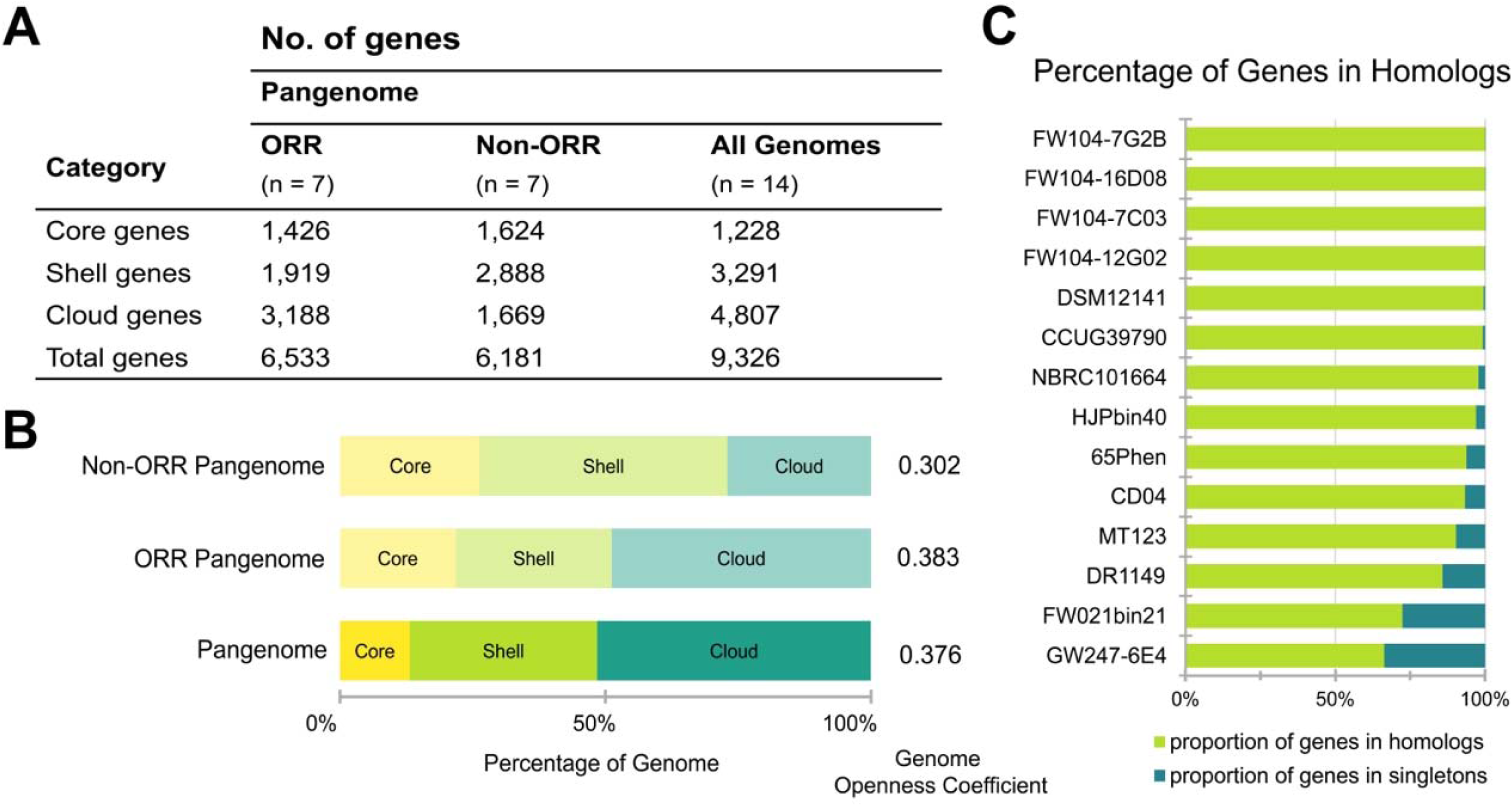
*Castellaniella* Pangenome. (**A**) The number of orthologs in each of the pangenome categories (core, shell, and cloud). The table columns represent the genomes included in the pangenome analysis. ORR signifies the pangenome was computed only using genomes derived from Oak Ridge Reservation whereas non-ORR included any publicly available *Castellaniella* sp. genomes that did not originate from this site. **(B)** The proportion of orthologs in each category with respect to the genomes included in the pangenome analysis. The core (yellow) category is made up of orthologs that are present in all genomes. The shell (green) category includes orthologs found in 16 ≤ x < 95% of genomes. The cloud (turquoise) category represents orthologs that are found in <16% of the genomes included in the analysis. The genome openness coefficient obtained for each of the pangenomes using a power law regression is indicated. **(C)** The distribution of homologs and singleton genes in the 14 *Castellaniella* sp. genomes.

To examine the importance of accessory genes in shaping the genomic diversity of *Castellaniella* species originating from different environments, we computed additional pangenomes for the seven genomes originating from ORR and for the remaining seven non-ORR *Castellaniella* genomes. Consistent with the greater taxonomic diversity of the ORR isolates, the ORR pangenome has a larger cloud genome (49% of the pangenome) than the non-ORR pangenome (27%). The *Castellaniella* pangenome (n=14) has a genome openness coefficient of 0.376 (Liao *et al*., 2021) **(Fig. 3B)**. The openness coefficient exists on a scale of 0 to 1. The closer the coefficient is to 1, the more open (i.e., has high potential gene pool growth). The degree of openness for the ORR pangenome (0.383) was larger than the value calculated for the *Castellaniella* pangenome and the value calculated for the non-ORR pangenome (0.302) **(Fig. 3B)**. These data suggest that *Castellaniella* originating from ORR have a greater propensity for acquiring genetic material and undergoing diversification events compared to their non-ORR counterparts. The heterogeneous nature (Smith *et al*., 2015) of stressors in the ORR subsurface may have promoted the higher level of genomic diversification observed among these strains (Chase *et al*., 2018). However, we note that a limitation of the comparison of the openness coefficient obtained for the *Castellaniella* pangenome (n=14) is the small number of high-quality *Castellaniella* genomes and MAGs available for inclusion in the analysis.

### Chromosomally Integrated Elements

Chromosomally integrated elements were found in all fourteen genomes with varying sizes and gene content, much of which is part of the *Castellaniella* cloud and shell genomes **(Table S5)**. The acquisition of these elements by HGT paired with their integration into the host’s genome is likely a factor contributing to the adaptation of these microorganisms to the unique environments at the ORR. On average, ORR genomes contained more chromosomally integrated elements than non-ORR genomes **(Fig. 4A)**. The chromosomally integrated elements most common in all *Castellaniella* genomes were genomic islands (GIs). Since the definition for GIs is broad, many elements which do not fit the criteria for other chromosomally integrated elements (e.g., integrative and conjugative elements, phages, insertion sequences) end up being classified as GIs (Langille *et al*., 2010). Non-ORR genomes had a greater proportion of GIs (49%) than ORR genomes (35%) **(Fig. 4A).** ORR genomes had more elements classified as integrative and conjugative elements (ICEs) and insertion sequences (IS) (14% and 4%, respectively) than did non-ORR genomes (10%).

**Figure 4.**
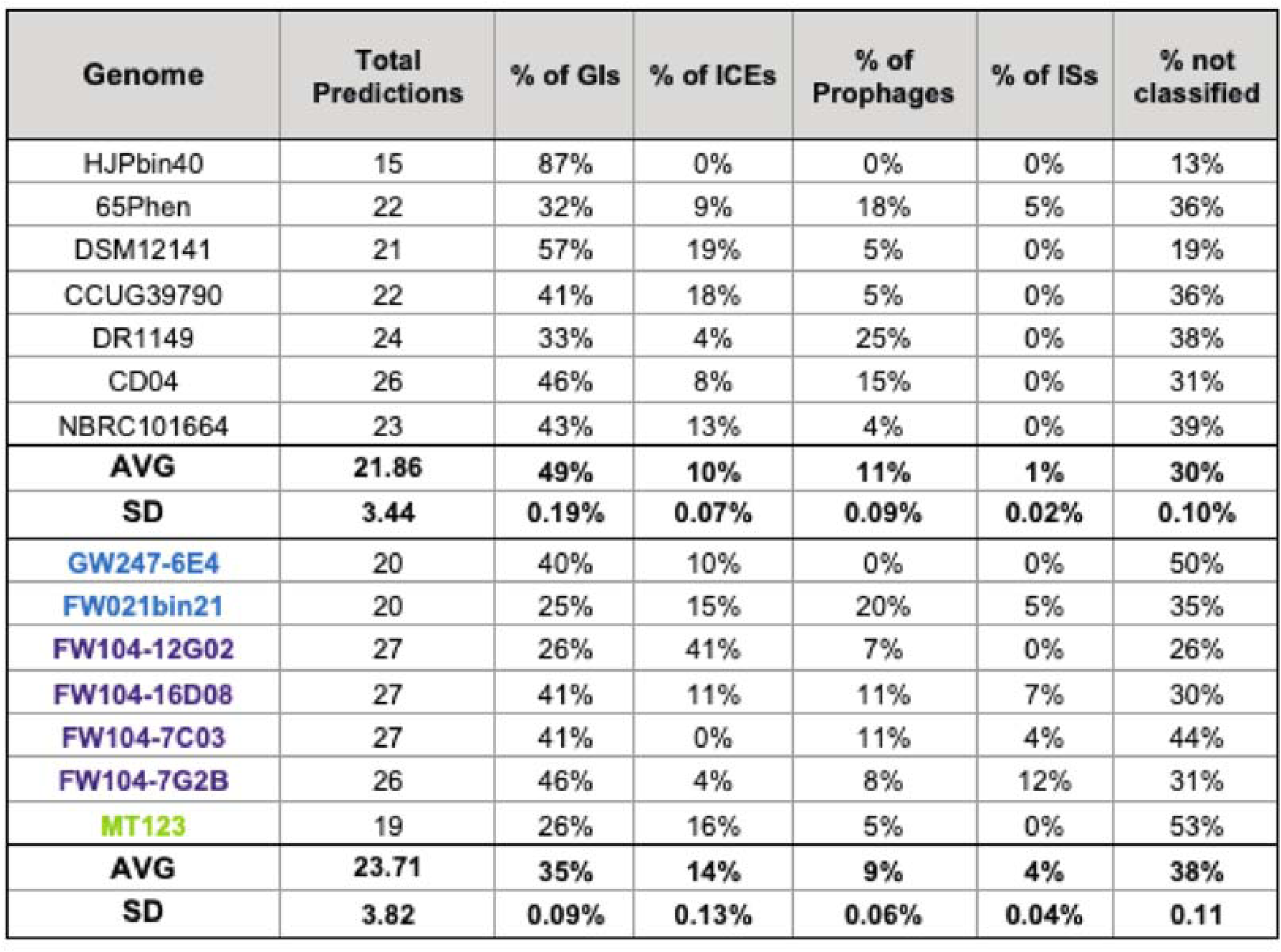
*Castellaniella* Chromosomally Integrated Elements. The total number of predictions made for each genome by IslandViewer4 as well as the percentage of these predictions further classified by additional annotation tools as being genomic islands (GIs), integrative and conjugative elements (ICEs), prophages or insertion sequences (ISs). The genomes are colored according to their matching ASV where purple represents ASV1 matching genomes, green represents the ASV2 matching genome and blue represents ORR genomes which do not match either ASV.

### Key functional traits for persistence in the ORR subsurface

Focusing on the strains representing *Castellaniella* ASV1 (FW104-16D08, FW104-12G02, FW104-7G2B, FW104-7C03) and ASV2 (MT123), we examined key functional traits that may facilitate their survival at the site. We sought to determine which of these traits were associated with the identified mobile elements and to what extent those elements may shape the fitness of the *Castellaniella* populations at this site.

### Acid tolerance

Bacterial populations in the ORR subsurface experience a wide range of pH, from highly acidic to neutral. In locations where the relative abundance of ASV1 and ASV2 is 0.5% or greater, both have similar environmentally defined pH ranges: ASV1 (3.9-7.4) and ASV2 (4.0-7.0). This is in line with prior studies which have found that pH preference *in situ* is deeply conserved in prokaryotic lineages (Martiny *et al*., 2015). The maximal relative abundance of both ASVs occurs in FW104 which has an average pH of 5.6. This observation is consistent with the pH optima for the ASV1-matching strains FW104-12G02 (pH 5.5), FW104-7C03 (pH 5.5), and FW104-7G2B (pH 5.5-7.0) **(Table S6)**. In contrast, ASV1-matching strain FW104-16D08 and ASV2-matching strain MT123 had higher pH optima, around pH 6-8 and pH 7-8, respectively. However, both strains still grew robustly at pH 5.5, with MT123 having its highest carrying capacity at pH 5.5, exceeding that of the ASV1 representatives at this pH. Only FW104-12G02, FW104-7G2B, and FW104-7C03 had significant growth at pH 5, and none of the strains had significant growth at pH < 5. In the field, the populations represented by ASV1 and ASV2 may persist in a dormant state < pH 5 and undergo blooms as pH increases, as was observed in the Spain *et al*. (2007) study when the subsurface pH was raised with bicarbonate. Additionally, our usage of batch cultures for determining pH tolerance may result in an underestimation of the pH range permissive towards growth (Goff *et al*., 2022b). Indeed, the permissive pH range for MT123 growth in laboratory batch culture extends down to 5.0 when grown aerobically without nitrate **(Fig. S7)**, eliminating accumulation of the protonophore nitrous acid (Sijbesma *et al*., 1996).

We examined acid tolerance systems of these key ORR isolates in the context of the larger *Castellaniella* pangenome **(Fig. S8)**. In general, *Castellaniella* genomes have a small repertoire of acid tolerance genes, in line with the observation that our isolates only tolerate weakly acidic conditions in the laboratory. The *Castellaniella* core genome includes the following genes known to be associated with low pH tolerance (but not exclusively involved in low pH tolerance): (1) RecA for DNA repair (Thompson & Blaser, 1995), (2) DnaK for protein re-folding (Tomoyasu *et al*., 2012), (3) the Pst phosphate transporter (Ryan *et al*., 2015), and (4) genes involved in the arginine-dependent low pH tolerance system (Ryan *et al*., 2015). MT123 encodes an additional aspartate-based (Hu *et al*., 2010) low pH tolerance system while FW104-12G02, FW104-7C03, FW104-7G2B, and FW104-16D08 encode a GI-associated glycerol-3-phosphate transporter (Ryan *et al*., 2015).

### Denitrification activity

Classically, *Castellaniella* are characterized as complete denitrifiers, reducing nitrate to dinitrogen (Kämpfer *et al*., 2006). Indeed, we observed that the ASV2-representative MT123 reduces nitrate to dinitrogen gas with no accumulation of nitrous oxide. MT123 NosZ was active in both the pH 7 and pH 5.5 cultures **(Table S7).** Based on bioinformatic analyses, the MT123 NosZ is a Clade II type (Hallin *et al*., 2018). Interestingly, in many denitrifiers, Clade II NosZ activity is eliminated below pH 6-6.5 (Carreira *et al*., 2020, Olaya-Abril *et al*., 2021). In our analysis, we also identified several variants on the *Castellaniella* denitrification pathway. For example, the ORR MAG, FW021bin21, lacks the nitrate reductase operon (*narGHJI*) as well as the nitrate/nitrite transporter gene (*narK*) (**Fig. 5A, B)**. Interestingly, the nitrous oxide genes for this MAG were found within a region predicted to be a genomic island. Six of the *Castellaniella* genomes have the NAD(P)H-dependent nitrite reductase genes (*nirBD*) **(Fig 5B)**. Out of those six genomes, only ORR genome MT123 contained the nitrite transporter gene *nirC*. Two of the non-ORR genomes, NBRC101664 and CD04, have a second copy of the nitrous oxide reductase operon (*nosXLYFDZR*) **(Fig 5B)**.

**Figure 5.**
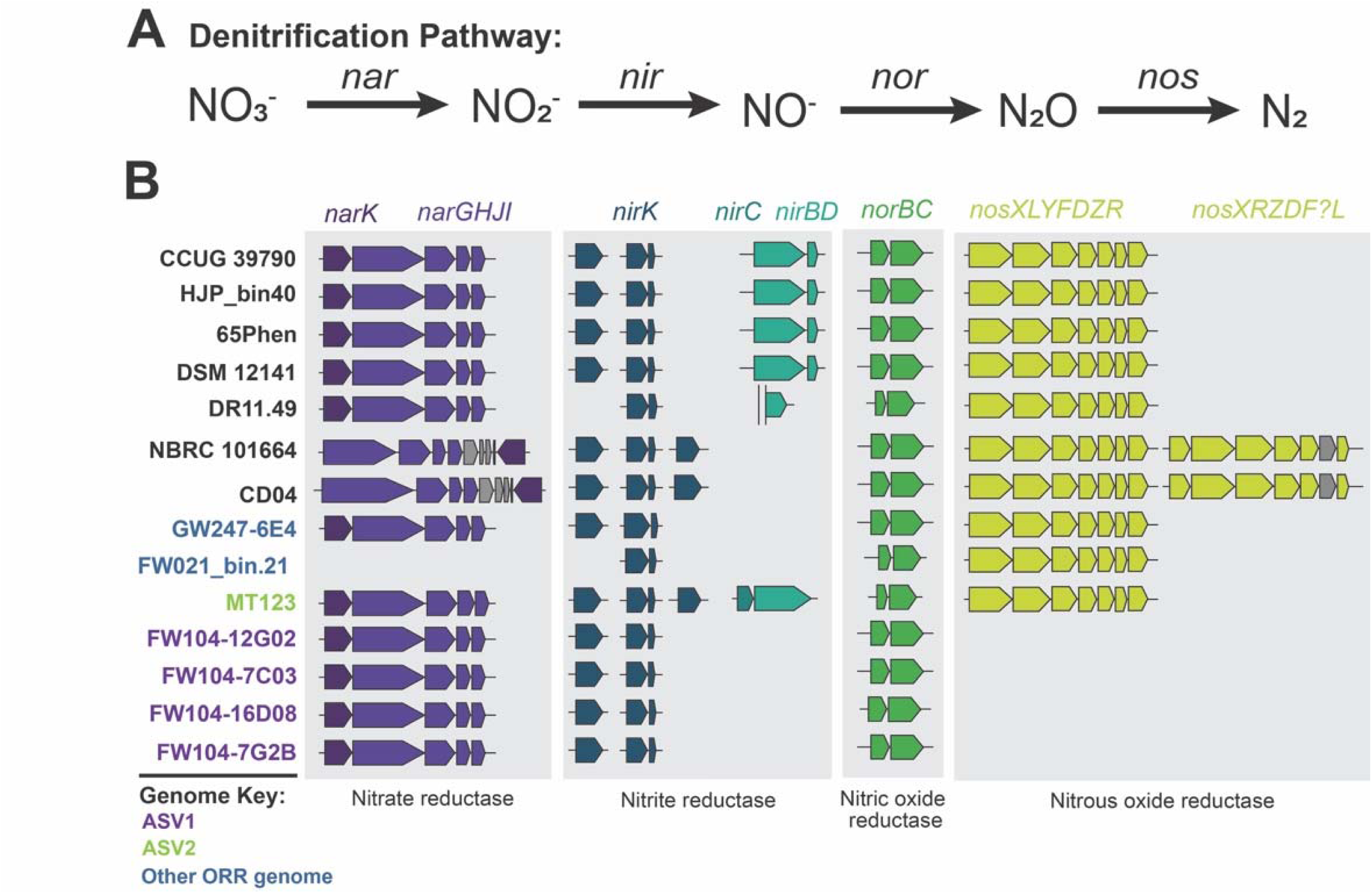
Denitrification Pathway. (**A**) Schematic representation of the denitrification pathway and the genes associated with each step. **(B)** The distribution of denitrification genes in the fourteen *Castellaniella* genomes. Non-ORR genomes are listed in black and ORR genomes are genomes are colored according to their matching ASV where purple represents ASV1 matching genomes, green represents the ASV2 matching genome and blue represents genomes which do not match either ASV. Genes shown include those encoding the nitrate/nitrite transporter (*narK*), the respiratory nitrate reductase and associated assembly proteins (*narGHJI*), the copper-containing respiratory nitrite reductase (*nirK*), the nitrite transporter (*nirC*), the NAD(P)H nitrite reductase (*nirBD*), the cytochrome c-dependent nitric oxide reductase (*norBC*), and the nitrous oxide reductase and associated assembly proteins (*nosXLYFDZR*).

Perhaps the most extreme instances of a denitrification pathway variants are the four ASV1-matching strains which lack the *nos* operon **(Fig 5B)**. The *nos* operon locus among *Castellaniella* (adjacent to an arginine tRNA gene) is highly conserved. Our analysis of this highly conserved region of the genome in ASV1-matching strains revealed a heavy metal homeostasis gene had been inserted at this location as part of a larger chromosomally integrated element. tRNA genes are common insertion sites for mobile genetic elements in microbial chromosomes (Boyd *et al*., 2009, Toleman & Walsh, 2011). The acquisition of a heavy metal homeostasis gene may contribute to the survival of these ASV1-matching strains within the subsurface at ORR. However, the absence of the *nos* operon in these four ASV1-matching strains suggests the final step in their dentification pathways is the production of the greenhouse gas nitrous oxide. Using strain FW104-7G2B as a representative, we confirmed the reduction of nitrate to nitrous oxide without further reduction to dinitrogen gas **(Table S7).** We propose that this inability to perform nitrous oxide reduction contributes to the apparent lower fitness of ASV1 at the site relative to ASV2 **(Fig 1B, C).** ASV1 strains are predicted to have lower molar growth yields from partial denitrification compared to complete denitrifiers (Koike & Hattori, 1975).

Additionally, nitrous oxide is cytotoxic; its accumulation can inactivate vitamin B12-dependent enzymes involved in methionine and DNA synthesis (Sullivan *et al*., 2013). ASV1 strains may be heavily dependent on nitrous-oxide-consuming neighbors to relieve this toxicity.

### Heavy metal homeostasis

There is emerging evidence to suggest that metals in combination interact synergistically or in an antagonistic manner within bacterial systems (Goff *et al*., 2023, Pormohammad *et al*., 2023). As the ORR subsurface is contaminated by complex and heterogenous mixtures of metals, we focused on microbial resistance to the metal composition present in the FW104 well **(Table S1),** as FW104 has the highest recorded *Castellaniella* relative abundance of all ORR sampling sites **(Fig. 1B,C)**. FW104 is also the origin of our ASV1 and ASV2 representative strains. The ASV2 representative MT123 experienced only a minor growth defect with the metal mixture, specifically, an extended lag phase. In contrast, the growth of the ASV1 representatives FW104-16D08, FW104-12G02, FW104-7G2B, FW104-7C03 was significantly inhibited by the FW104 metal mixture (**Fig. S9**). These findings add context to the observation that ASV2 is at a greater abundance in the FW104 groundwater (and across all samples) relative to ASV1 **(Fig. 1B, C).** Metal tolerance may be a significant distinguishing factor between co-existing *Castellaniella* ecotypes in the ORR subsurface.

We examined heavy metal homeostasis gene (HMHG) content in *Castellaniella* genomes: encompassing genes involved in the import, export, intracellular trafficking, and transformation of metals (Chandrangsu *et al*., 2017). ORR genomes had significantly higher (Student’s two-sided t-test *p < 0.05*) (**Fig. 6A**) HMHG counts than non-ORR genomes. ORR genomes encode an average of 90 HMHGs compared to 80 in non-ORR genomes. While many of the non-ORR genomes do originate from anthropogenically-impacted sites like wastewater treatment plants (Foss *et al*., 1998, Woo, 2017), heavy metal concentrations in these areas are expected to be lower than the ORR site. Strain 65Phen had the lowest HMHG count of all *Castellaniella*, reflective of its origin from unimpacted soils (Foss *et al*., 1998). Among the ORR genomes, the strain GW247-6E4 genome had the lowest HMHG count. Notably, this strain is not reflected in any of the ORR *Castellaniella* ASVs, suggestive of its low relative abundance at the site.

**Figure 6.**
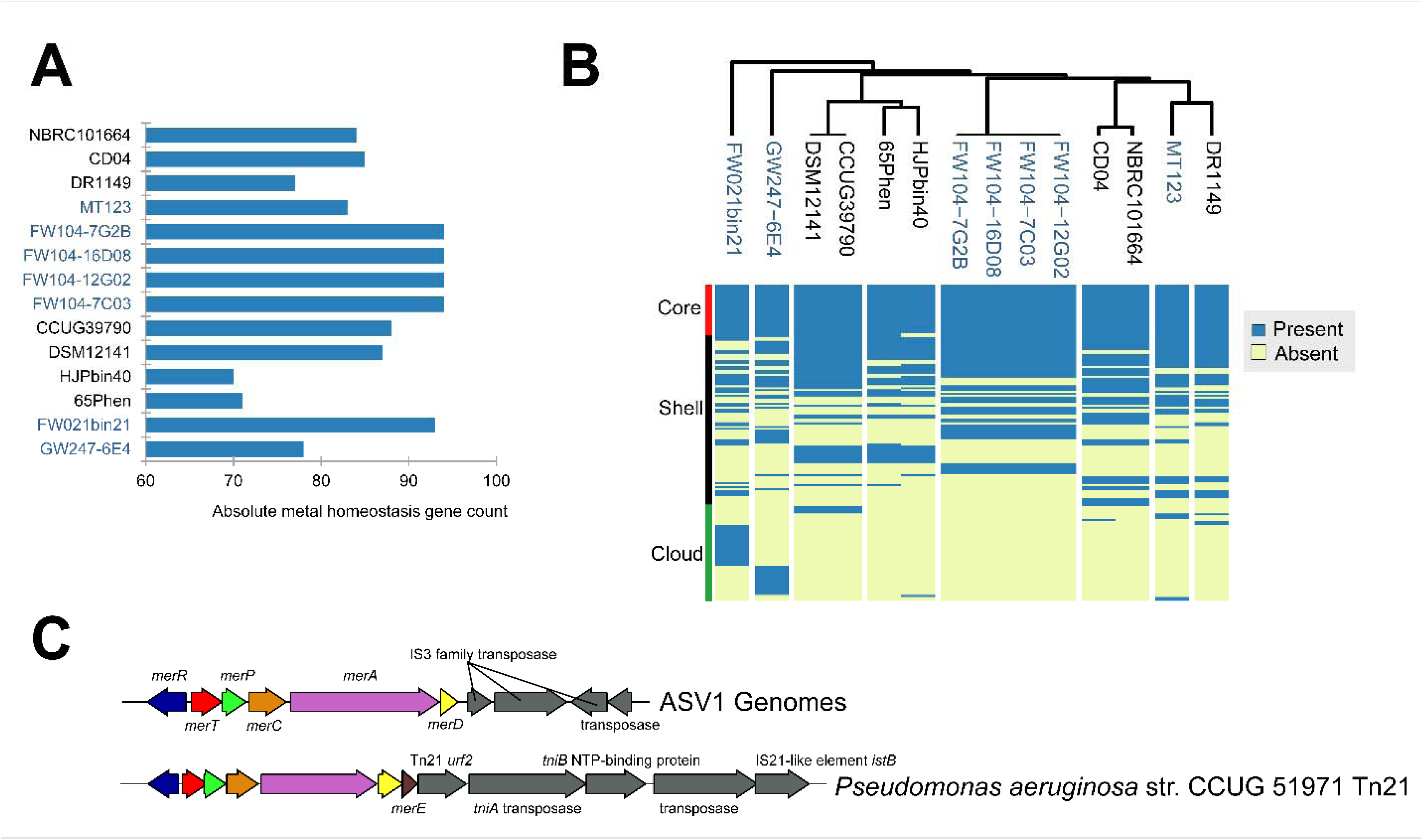
Metal homeostasis genes. (**A**) Counts of heavy metal homeostasis genes (HMHG) identified in *Castellaniella* genomes. **(B)** Presence/absence matrix of individual HMHGs in *Castellaniella* genomes. Whether a particular gene belongs to the core, shell, or cloud genome is indicated. Clustering was performed using a Euclidian distance metric **(C)** Mercury transposon observed in the ASV1-representative genomes compared to a canonical *Tn*21 mercury transposon in *P. aeruginosa*.

However, the total number of HMHGs did not correlate with their degree of metal mixture tolerance among our ASV1 and ASV2-representative genomes **(Fig. S9).** Instead, we propose that the presence or absence of specific HMHGs drives the differences in metal tolerance between these two ORR ecotypes. The MT123 genome encodes 17 HMHGs not found in any of the ASV1 representative genomes **(Table S8)**. These include two P-type metal ion-pumping ATPases and several copper and mercury resistance genes. Most of these ASV2-specific genes are found on a GI. Acquisition of this GI may have increased the ASV2 population fitness within this niche relative to the ASV1 population.

Cloud and shell HMHGs are frequently associated with genomic islands (Rodriguez-Valera *et al*., 2016). Within the ORR genomes, 0-40% (median = 16%) of the identified HMHGs are located on genomic islands. These are similar values to what was previously observed for *Rhodanobacter* from the ORR subsurface (Peng *et al*., 2022). For example, the copper resistance genes inserted at the location of the *nos* operson of the ASV1 representative genomes are part of a larger predicted genomic island. We also identified a mercury resistance transposon containing the *merRTPCAD* operon in these ASV1 representative genomes **(Fig. 6B).** This *mer* operon resembles the well-characterized gram-negative Tn*21,* Tn*501,* and pKLH2 mercuric ion resistance operons (Osborn *et al*., 1997). Tn*21* and Tn*501* were both shown to be highly abundant in the mercury-contaminated New Hope Pond site near the former S-3 ponds (Barkay *et al*., 1991). Mercury is also part of the S-3 contaminant profile (Revil *et al*., 2013,

Kothari *et al*., 2019). Overall, the accessory HMHG profiles are consistent with the phylogenomic tree shown in **Figure 2A**, with more closely related genomes having more similar shell and cloud HMHG content, suggesting that these traits are conserved on a very fine-grained level (**Fig. 6C**). This is consistent with recent findings that host phylogeny significantly controls the acquisition of functional traits like resistance genes (Finks & Martiny, 2023).

### Global Significance

The majority of sequenced *Castellaniella* genomes analyzed in this study originate from anthropogenically-impacted sites like the ORR subsurface and anaerobic digesters. We downloaded an additional 898 *Castellaniella* 16S rRNA gene sequences from the SILVA database **(Fig. 7A, Table S8).** Of these, 680 (76%) originate from globally distributed anthropogenically impacted sites **(Fig. 7B)** that we broadly categorized as “Terrestrial-Contaminated”, “Aquatic-Contaminated”, “Solid Waste”, “Built Environment”, and “Wastewater”. We propose that this genus is specialized for growth in sites which are typically impacted by multiple simultaneous stressors (i.e., heavy metals, organics, nitrogen pollution, and/or low pH). This is likely due, in part, to the ability of members of this genus to acquire novel genes via HGT. For example, in addition to HMRGs, many of the genomic islands observed in the *Castellaniella* genomes (both ORR and non-ORR) carry genes involved in acid tolerance, aromatic carbon degradation, toxin/antitoxin defense systems, and biofilm formation.

**Figure 7.**
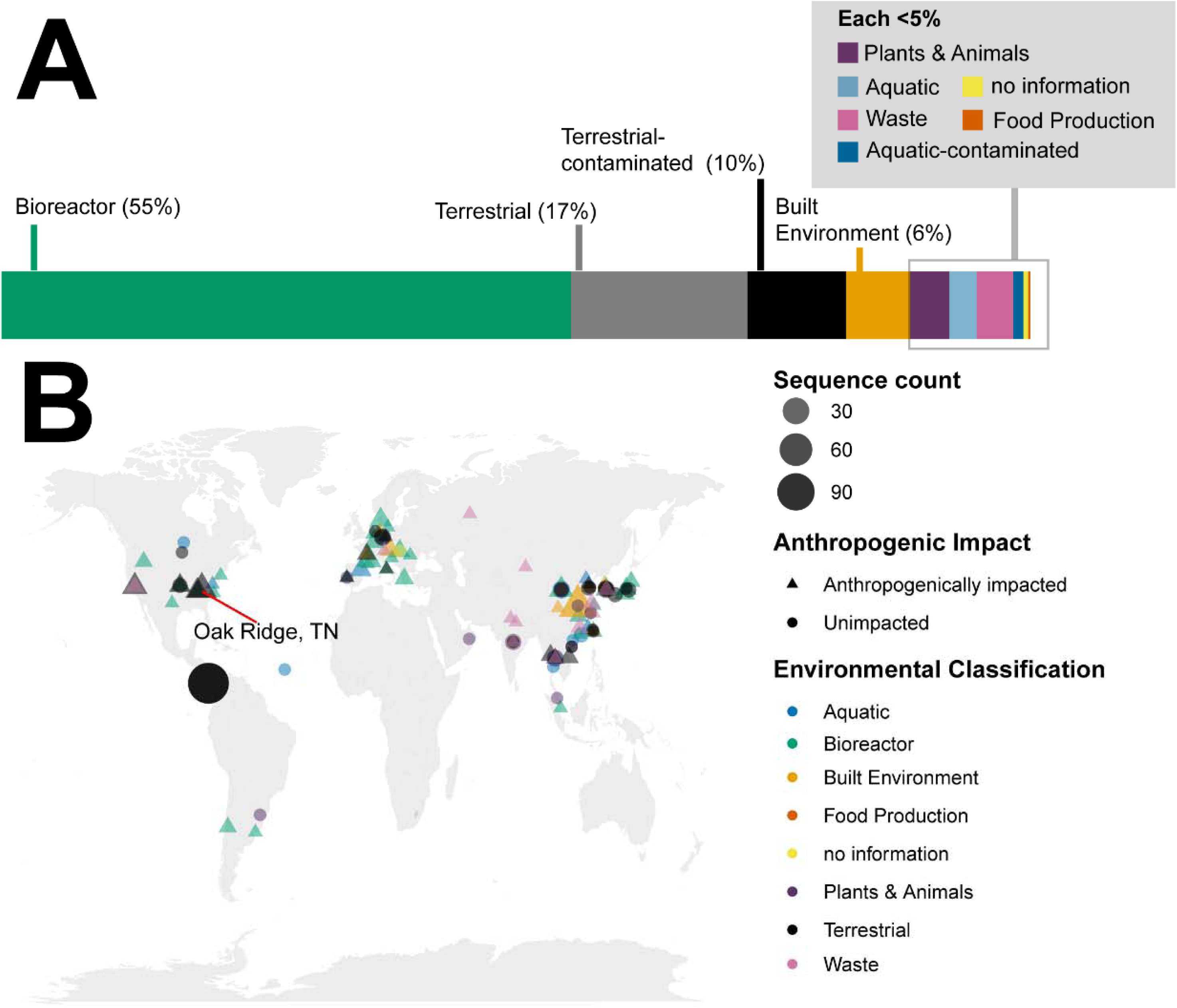
Global distribution of *Castellaniella* 16S rRNA gene sequences. *Castellaniella* 16S rRNA gene sequences were retrieved from the SILVA database (v.138.1). (A) The bar graph shows the percentages of 16S rRNA gene sequences (n = 898 total) associated with each environment classification. (B) The map shows the geographic distribution of the 16S rRNA gene sequences. Colors indicate the environmental classification. Shapes indicated sites that are anthropogenically impacted (triangle) or unimpacted (circle). Shapes are scaled to reflect the count of sequences originating from a single location.

## CONCLUSIONS

The greater presence of chromosomally integrated elements found throughout ORR *Castellaniella* genomes relative to non-ORR genomes suggest that these extant lineages have a high propensity to acquire and integrate novel genetic material into their genomes.

Investigation of the representative functions for genes encoded within these chromosomally integrated regions in the ORR genomes revealed numerous acid tolerance, denitrification, and heavy metal homeostasis genes. Paired with the fact that *Castellaniella* have been shown to persist in the subsurface at ORR, these data demonstrate the importance of horizontal gene transfer in the diversification and adaptation of microorganisms in this multi-stressor environment. The discussion of niche-specific adaptations raises the question of whether contaminants found in the subsurface prompted these differentiation events through increased rates of horizontal gene transfer (Chen *et al*., 2023) or whether the isolates already harbored the functionalities necessary to allow for their survival in such an environment (Hemme *et al*., 2016). The answer likely falls somewhere between those two scenarios. On-going mutant fitness and omics-based analyses of these ORR *Castellaniella* isolates seek to further characterize the genomic controls of *Castellaniella* persistence at ORR.

## DECLARATIONS

### Ethics approval and consent to participate

Not applicable

### Consent for publication

Not applicable

### Availability of data and material

Assembled genomes and annotations can be found in the KBase narrative: https://doi.org/10.25982/135835.33/2203552 (for review: https://kbase.us/n/135835/33/). The narrative also contains the output files for the pangenome analysis.

### Authors’ contributions

Study conception and design were carried out by JLG and EGS. Experimental work was performed by EGS, JLG, KLD, KAH, JVK, JH, and MPT. Data analyses and figure generation were performed by EGS, JLG, and KLD. Genome sequencing and assembly were performed by LML, TNN, JVK, and J-MC. The first draft of the manuscript was written by EGS and JLG. All authors were involved in revisions of subsequent versions of the manuscripts. DAS, RC, APA, AMD, and MWWA oversaw the research and acquired funding.

## Acknowledgements

Not applicable

