## Supplemental Materials for "Genomic and environmental controls on *Castellaniella* biogeography in an anthropogenically disturbed subsurface"

**Table S1. FW104 COMM Composition.**

| **Metal** | **FW104 Concentration (µM)**^1^ | **COMM Concentration (µM)** |
| --- | --- | --- |
| Al | 215 | 215 |
| U | 78 | 80 |
| Mn | 3,020 | 3,000 |
| Ni | 19 | 20 |
| Co | 1.68 | 2 |
| Fe | 9.82 | 10 |
| Cu | 0.567 | 1 |
| Cd | 0.944 | 1 |

^1^Metal concentrations from Thorgersen et al. (2019)

**Table S2. Oak Ridge Reservation (ORR) *Castellaniella* 16S V4 Region ASVs**

| ASV | Sequence |
| --- | --- |
| 1 | TACGTAGGGTGCAAGCGTTAATCGGAATTACTGGGCGTAAAGCGTGCGCAGGCGGTTCGGAAAGAAAGGTGTGAAATCCCAGGGCTTAACCTTGGAACTGCACTTTTAACTACCGGGCTAGAGTACGTCAGAGGGGGGTAGAATTCCACGTGTAGCAGTGAAATGCGTAGAGATGTGGAGGAATACCGATGGCGAAGGCAGCCCCCTGGGATGATACTGACGCTCATGCACGAAAGCGTGGGGAGCAAACAGG |
| 2 | TACGTAGGGTGCAAGCGTTAATCGGAATTACTGGGCGTAAAGCGTGCGCAGGCGGTTCGGAAAGAAAGGTGTGAAATCCCAGGGCTTAACCTTGGAACTGCACTTTTAACTCCCGAGCTAGAGTACGTCAGAGGGGGGTAGAATTCCACGTGTAGCAGTGAAATGCGTAGAGATGTGGAGGAATACCGATGGCGAAGGCAGCCCCCTGGGATGATACTGACGCTCATGCACGAAAGCGTGGGGAGCAAACAGG |
| 3 | TACGTAGGGTGCAAGCGTTAATCGGAATTACTGGGCGTAAAGCGTGCGCAGGCGGTTCGGAAAGAAAGGTGTGAAATCCCAGGGCTTAACCCTGGAACTGCACTTTTAACTCCCGAGCTAGAGTACGTCAGAGGGGGGTAGAATTCCACGTGTAGCAGTGAAATGCGTAGAGATGTGGAGGAATACCGATGGCGAAGGCAGCCCCCTGGGATGATACTGACGCTCATGCACGAAAGCGTGGGGAGCAAACAGG |
| 4 | TACGTAGGGTGCAAGCGTTAATCGGAATTACTGGGCGTAAAGCGTGCGCAGGCGGTTCGGAAAGAAAGGTGTGAAATCCCAGGGCTTAACCTTGGAACTGCACTTTTAACTACCGGGCTAGAGTACGTCAGAGGGGGGTAGAATTCCACGTGTAGCAGTGAAATGCGTAGAGATGTGGAGGAATACCGATGGCGAAGGCAGCCCCCTGGGATGATACTGACGCTCATGCACGAAAGCGTGGGGAGCAAGCAGG |
| 5 | TACGTAGGGTGCGAGCGTTAATCGGAATTACTGGGCGTAAAGCGTGCGCAGGCGGTTCGGAAAGAGGGGTGTGAAATCCCGGGGCTTAACCCCGGAACTGCACTTCTAACTACCGGGCTAGAGTACGTCAGAGGGGGGTAGAATTCCACGTGTAGCAGTGAAATGCGTAGAGATGTGGAGGAATACCGATGGCGAAGGCAGCCCCCTGGGATGATACTGACGCTCAGGCACGAAAGCGTGGGGAGCAAACAGG |
| 6 | TACGTAGGGTGCAAGCGTTAATCGGAATTACTGGGCGTAAAGCGTGCGCAGGCGGTTCGGAAAGAGAGGTGTGAAATCCCAGGGCTTAACCCTGGAACTGCACTTCTAACTACCGGGCTAGAGTACGTCAGAGGGGGGTAGAATTCCACGTGTAGCAGTGAAATGCGTAGAGATGTGGAGGAATACCGATGGCGAAGGCAGCCCCCTGGGACGATACTGACGCTCAGGCACGAAAGCGTGGGGAGCAAACAGG |
| 7 | TACGTAGGGTGCAAGCGTTAATCGGAATTACTGGGCGTAAAGCGTGCGCAGGCGGTTCGGAAAGAGAGGTGTGAAATCCCAGGGCTCAACCCTGGAACTGCACTTCTAACTACCGGGCTAGAGTACGTCAGAGGGGGGTAGAATTCCACGTGTAGCAGTGAAATGCGTAGAGATGTGGAGGAATACCGATGGCGAAGGCAGCCCCCTGGGATGATACTGACGCTCATGCACGAAAGCGTGGGGAGCAAACAGG |
| 8 | TACGTAGGGTGCAAGCGTTAATCGGAATTACTGGGCGTAAAGCGTGCGCAGGCGGTTCGGAAAGAGGGGTGTGAAATCCCAGGGCTTAACCCTGGAACTGCACTCCTAACTACCGGGCTAGAGTACGTCAGAGGGGGGTAGAATTCCACGTGTAGCAGTGAAATGCGTAGAGATGTGGAGGAATACCGATGGCGAAGGCAGCCCCCTGGGATGATACTGACGCTCATGCACGAAAGCGTGGGGAGCAAACAGG |
| 9 | TACGTAGGGTGCAAGCGTTAATCGGAATTACTGGGCGTAAAGCGTGCGCAGGCGGTTCGGAAAGAAAGGTGTGAAATCCCGGGGCTTAACCTCGGAACTGCACTTTTAACTACCGGGCTAGAGTACGTCAGAGGGGGGTAGAATTCCACGTGTAGCAGTGAAATGCGTAGAGATGTGGAGGAATACCGATGGCGAAGGCAGCCCCCTGGGATGATACTGACGCTCATGCACGAAAGCGTGGGGAGCAAACAGG |
| 10 | TACGTAGGGTGCGAGCGTTAATCGGAATTACTGGGCGTAAAGCGTGCGCAGGCGGTTCGGAAAGAGGGGTGTGAAATCCCGGGGCTTAACCCCGGAACTGCACTCCTAACTACCGGGCTAGAGTACGTCAGAGGGGGGTAGAATTCCACGTGTAGCAGTGAAATGCGTAGAGATGTGGAGGAATACCGATGGCGAAGGCAGCCCCCTGGGATGATACTGACGCTCAGGCACGAAAGCGTGGGGAGCAAACAGG |
| 11 | TACGTAGGGTGCAAGCGTTAATCGGAATTACTGGGCGTAAAGCGTGCGCAGGCGGTTCGGCAAGAAAGGTGTGAAATCCCAGGGCTTAACCTTGGAACTGCACTTTTAACTACCGGGCTAGAGTACGTCAGAGGGGGGTAGAATTCCACGTGTAGCAGTGAAATGCGTAGAGATGTGGAGGAATACCGATGGCGAAGGCAGCCCCCTGGGATGATACTGACGCTCATGCACGAAAGCGTGGGGAGCAAACAGG |
| 12 | TACGTAGGGTGCAAGCGTTAATCGGAATTACTGGGCGTAAAGCGTGCGCAGGCGGTTCGGAAAGAGAGGTGTGAAATCCCAGGGCTTAACCTTGGAACTGCACTTTTAACTACCGGGCTAGAGTACGTCAGAGGGGGGTAGAATTCCACGTGTAGCAGTGAAATGCGTAGAGATGTGGAGGAATACCGATGGCGAAGGCAGCCCCCTGGGATGATACTGACGCTCATGCACGAAAGCGTGGGGAGCAAACAGG |

**Table S3. COGs used for phylogenomic tree**

| COG ID | Protein Product | COG ID | Protein Product |
| --- | --- | --- | --- |
| COG0012 | COG0012 | COG0097 | RplF |
| COG0013 | AlaS | COG0098 | RpsE |
| COG0016 | PheS | COG0099 | RpsM |
| COG0018 | ArgS | COG0100 | RpsK |
| COG0030 | KsgA | COG0102 | RplM |
| COG0041 | PurE | COG0103 | RpsI |
| COG0046 | PurL | COG0105 | Ndk |
| COG0048 | RpsL | COG0126 | Pgk |
| COG0049 | RpsG | COG0127 | COG0127 |
| COG0051 | RpsJ | COG0130 | TruB |
| COG0052 | RpsB | COG0150 | PurM |
| COG0072 | PheT | COG0151 | PurD |
| COG0080 | RplK | COG0164 | RnhB |
| COG0081 | RplA | COG0172 | SerS |
| COG0082 | AroC | COG0185 | RpsS |
| COG0086 | RpoC | COG0186 | RpsQ |
| COG0087 | RplC | COG0215 | CysS |
| COG0088 | RplD | COG0244 | RplJ |
| COG0089 | RplW | COG0256 | RplR |
| COG0090 | RplB | COG0343 | Tgt |
| COG0091 | RplV | COG0504 | PyrG |
| COG0092 | RpsC | COG0519 | GuaA |
| COG0093 | RplN | COG0532 | InfB |
| COG0094 | RplE | COG0533 | QRI7 |
| COG0096 | RpsH |  |  |

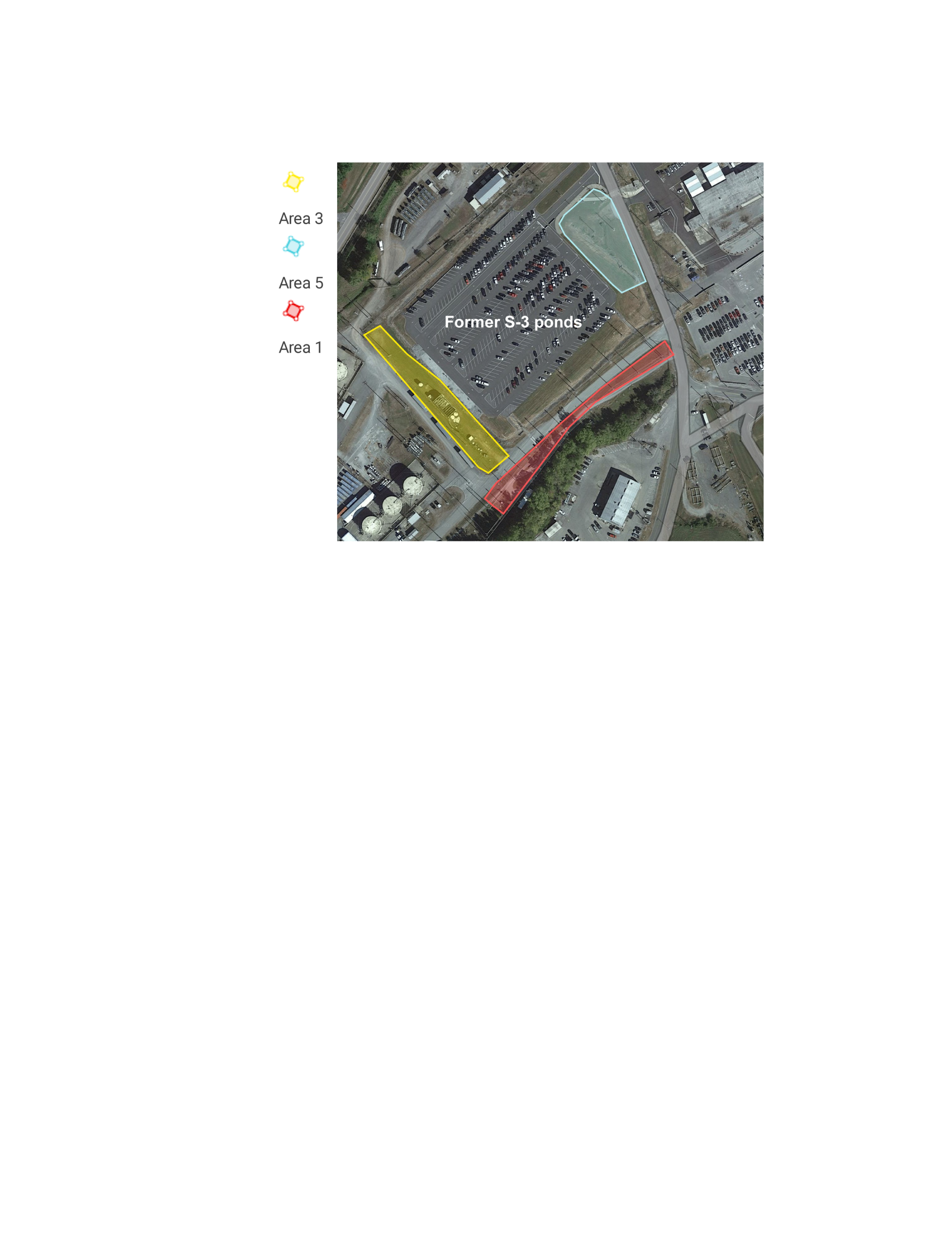
**Figure S1. ORR site map near the former S-3 ponds.** Sampling areas “3”, “5”, and “1” are marked on the map.

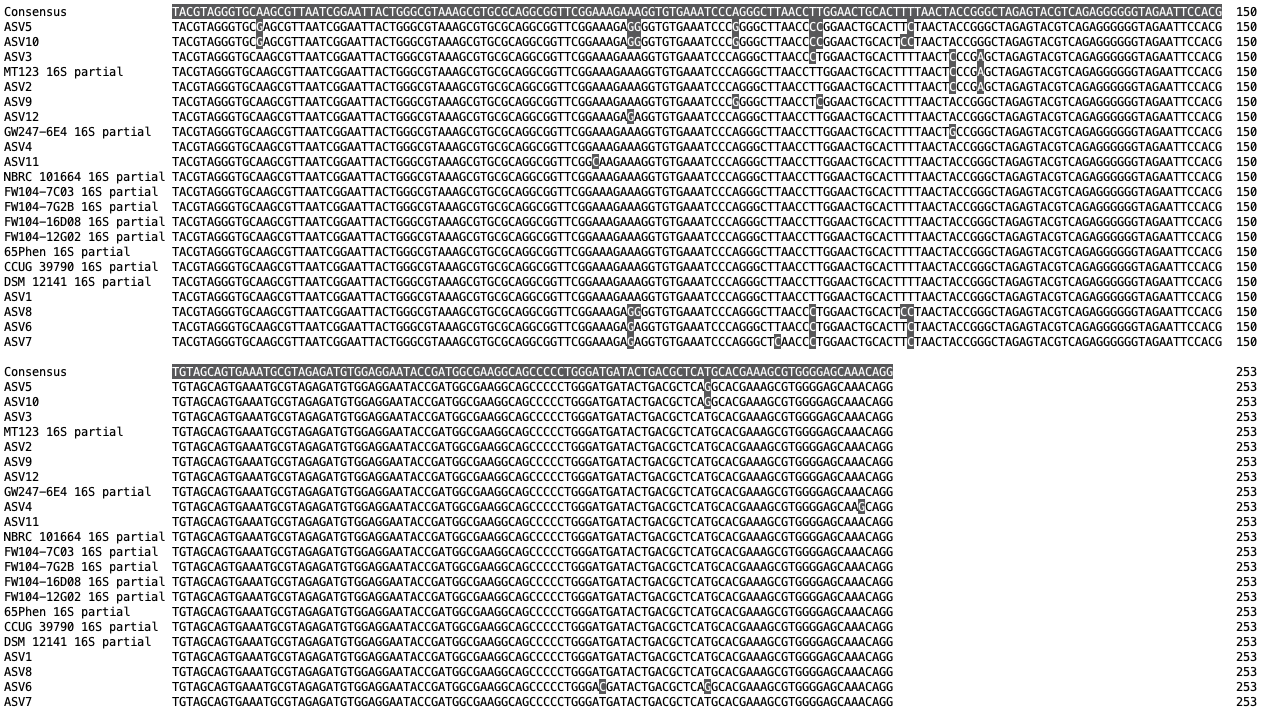

**Figure S2.** Alignment of Oak Ridge Reservation (ORR) *Castellaniella* 16S V4 region ASVs and partial 16S sequences from *Castellaniella* genomes. This alignment was used to generate the tree in **Figure 1A**.

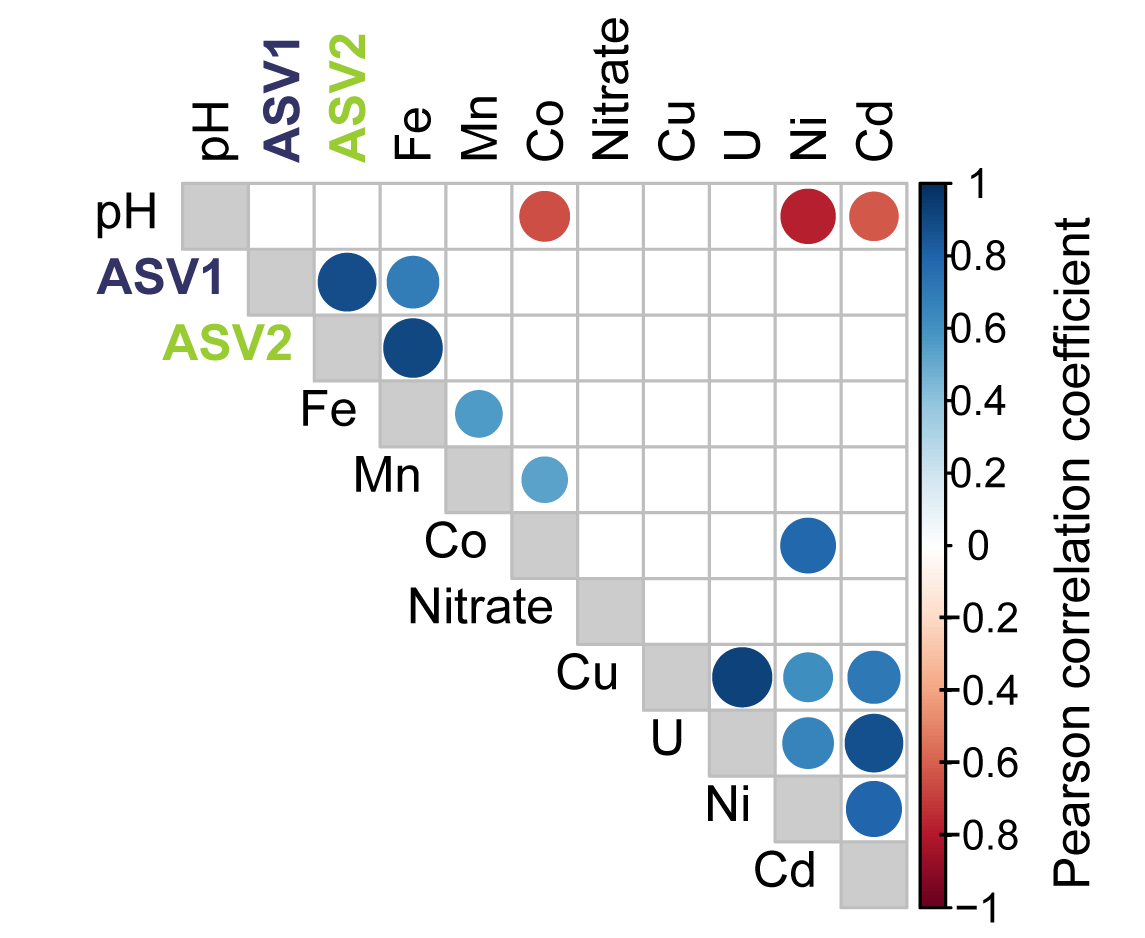

**Figure S3. Pearson correlation analysis.** Analysis incudes relative abundances of ASV1 and ASV2 as well as corresponding well geochemistry. Only statistically significant correlations are displayed (p <0.05). Circle colors reflect correlation coefficients. The correlation matrix was generated in R using the *corrplot* package (v0.92).

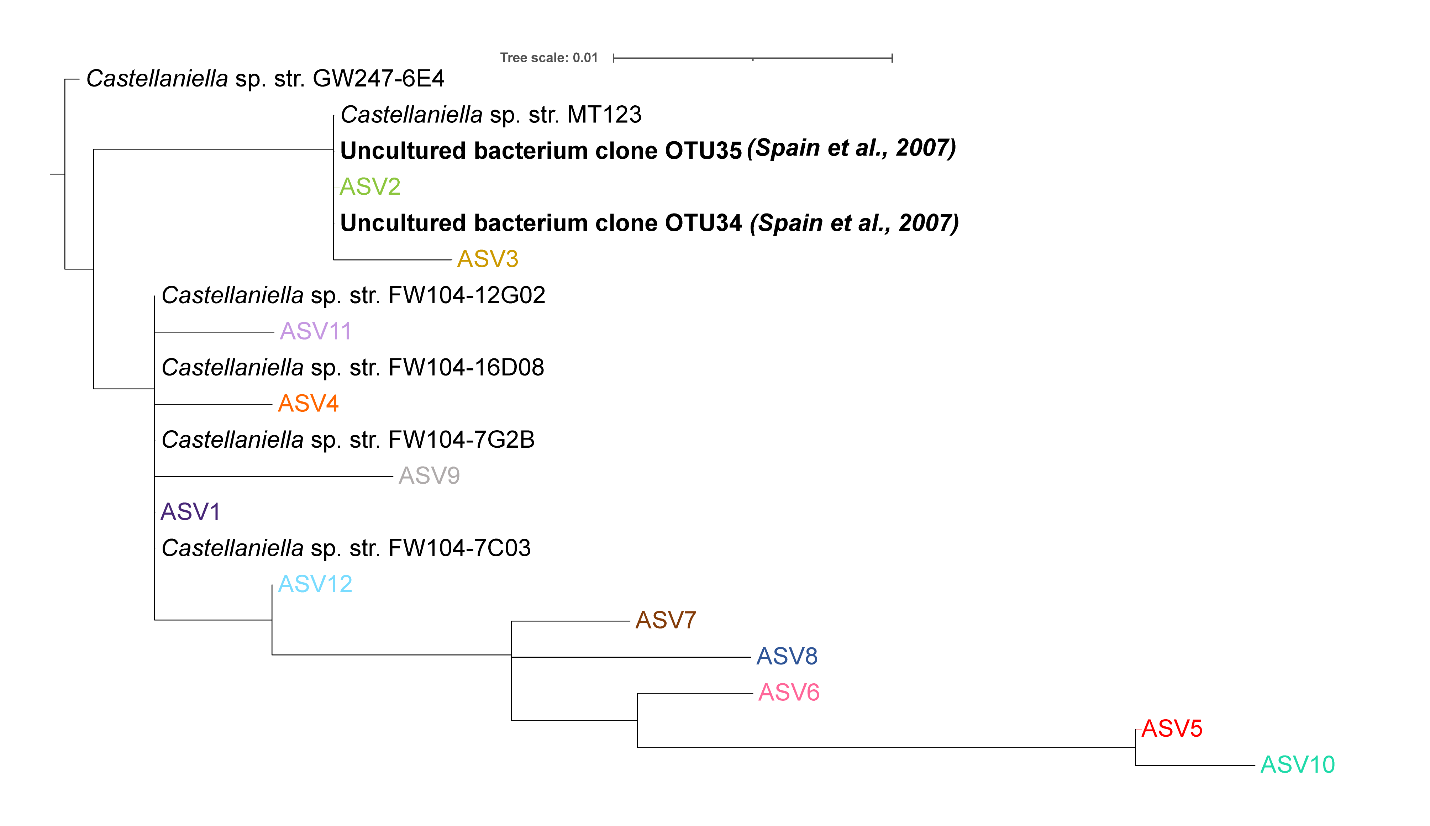

**Figure S4. Phylogenetic tree using partial 16S (V4) region sequences.** This tree includes sequenced ORR *Castellaniella* strains, ORR *Castellaniella* ASVs, and two uncultured bacterial clones from Spain et al. (2007).

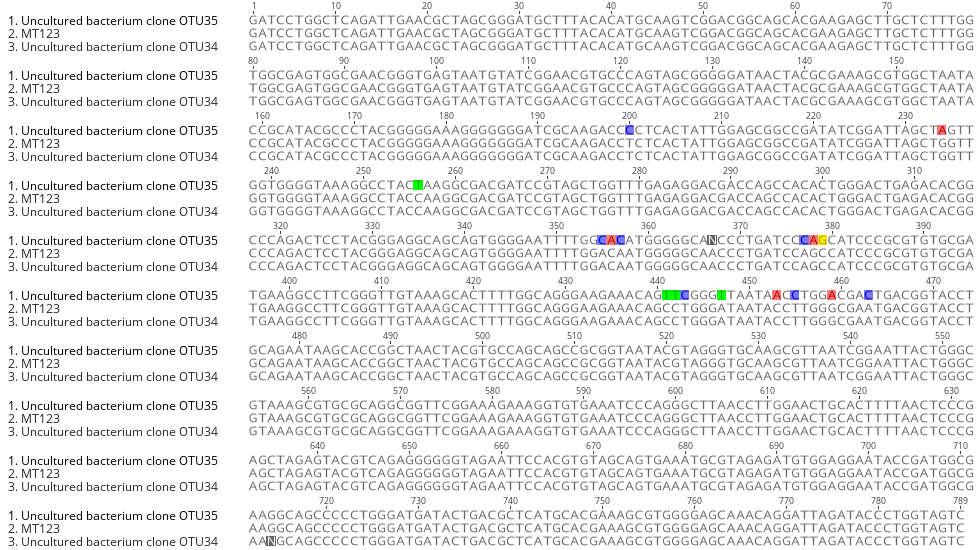

**Figure S5. Alignment of MT123 partial 16S sequence with *Castellaniella* clones from Spain et al. (2007).** The alignment was performed with MUSCLE v3.8.425 implemented in and visualized with Geneious Prime (v2022.0.2).

**Table S4. Basic Characteristics of Castellaniella genomes**. Publicly available genomes from non-ORR sites are unshaded. ORR genomes are shaded in grey. MAG: Metagenome assembled genome; Y: Yes; N: No

| **Strain** | **NCBI Classification** | **GTDBtk Classification** | **Accession Number** | **GOLD Ecosystem Category** | **Habitat Details** | **Contigs** | **Bases** | **GC%** | **CDS** | **MAG** |
| --- | --- | --- | --- | --- | --- | --- | --- | --- | --- | --- |
| HJP_bin40 | *C. defragrans* | *C. defragrans* | GCA_017848875.1 | Wastewater | Sludge from phenol-fed wastewater aerobic bioreactor | 48 | 3596155 | 69.47 | 3331 | Y |
| DSM 12141 | *C. defragrans* | *C. defragrans* | GCA_014203015.1 | Wastewater | Activated sludge | 45 | 3942929 | 69.20 | 3696 | N |
| CCUG 39790 | *C. defragrans* | *C. defragrans* | GCA_008801975.1 | Wastewater | Activated sludge | 345 | 3962237 | 69.14 | 3989 | N |
| NBRC 101664 | *C. caeni* | *C. caeni* | GCF_001592225.1 | Wastewater | Activated sludge | 60 | 3282217 | 64.99 | 3094 | N |
| DR_1_1.49 | *C. sp.* | *C. sp.* | GCA_019104865.1 | Wastewater | Anaerobic fermentation tank | 102 | 3463312 | 63.47 | 3475 | Y |
| CD04 | *C. caeni* | *C. caeni* | GCA_002894315.1 | Wastewater | Sludge from N_2_O producing bioreactor | 94 | 3414267 | 64.89 | 3297 | N |
| 65Phen | *C. defragrans* | *C. defragrans* | GCF_000612685.1 | Terrestrial | Ditch in a forest | 1 | 3952818 | 68.94 | 3616 | N |
| FW021_bin.21 | *C. sp.* | *C. sp.* | GCA_004321985.1 | Terrestrial | Groundwater from radionuclide waste site | 86 | 3407388 | 62.66 | 3384 | Y |
| MT123 | *C. defragrans* | *C. sp.* | Pending (available for review in associated KBase narrative) | Terrestrial | Groundwater from radionuclide waste site | 1 | 3366678 | 63.47 | 3202 | N |
| FW104-7C03 |  | *C. sp.* | Pending (available for review in associated KBase narrative) | Terrestrial | Groundwater from radionuclide waste site | 36 | 3105476 | 59.10 | 3004 | N |
| FW104-12G02 |  | *C. sp.* | Pending (available for review in associated KBase narrative) | Terrestrial | Groundwater from radionuclide waste site | 37 | 3105632 | 59.10 | 3005 | N |
| FW104-16D08 |  | *C. sp.* | Pending (available for review in associated KBase narrative) | Terrestrial | Groundwater from radionuclide waste site | 38 | 3105371 | 59.10 | 3003 | N |
| FW1047G2B |  | *C. sp.* | Pending (available for review in associated KBase narrative) | Terrestrial | Groundwater from radionuclide waste site | 40 | 3105685 | 59.10 | 3003 | N |
| GW247-6E4 |  | *C. sp.* | Pending (available for review in associated KBase narrative) | Terrestrial | Groundwater from radionuclide waste site | 59 | 3572056 | 64.65 | 3424 | N |

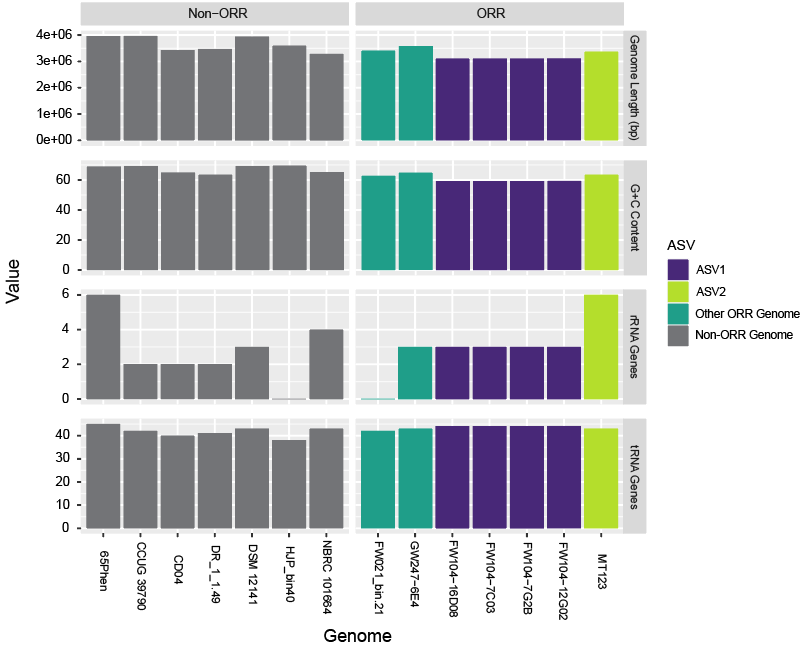

**Figure S6. General Genome Features.** This figure separates the genomes by origin (non-ORR and ORR) and presents bars which are colored based on the genome’s matching ASV. Genomes which match ASV1 are colored purple, genomes which match ASV2 are colored in green, and genomes which do not match either ASV are colored turquoise. The values of the bar represent the size of the genome in base pairs, percentage of G+C content, number of rRNA genes and number of tRNA genes found in each of the genomes.

OD600

| Time (h) | pH 4 | pH 4.5 | pH 5 | pH 5.5 | pH 6 | pH 7 | pH 8 |
| --- | --- | --- | --- | --- | --- | --- | --- |
| 0 | 0 | 0 | 0 | 0 | 0 | 0 | 0 |
| 1 | 0.04328 | 0.0222 | 0.0258333 | 0.03212 | 0.03352 | 0.0222 | 0.03166 |
| 2 | 0.05296 | 0.0328 | 0.0389333 | 0.04606 | 0.05156 | 0.03248 | 0.03124 |
| 3 | 0.05744 | 0.0347 | 0.0478333 | 0.04794 | 0.05834 | 0.02802 | 0.03516 |
| 4 | 0.05836 | 0.034075 | 0.0438 | 0.0595 | 0.06 | 0.02676 | 0.03476 |
| 5 | 0.06024 | 0.034475 | 0.0422 | 0.07718 | 0.06222 | 0.03234 | 0.03244 |
| 6 | 0.07044 | 0.034975 | 0.0417 | 0.08278 | 0.08044 | 0.03846 | 0.04498 |
| 7 | 0.0664 | 0.03465 | 0.0414667 | 0.10454 | 0.1001 | 0.04698 | 0.04794 |
| 8 | 0.07408 | 0.0356 | 0.0413333 | 0.12922 | 0.1231 | 0.05382 | 0.05058 |
| 9 | 0.07222 | 0.037075 | 0.0408 | 0.16162 | 0.1431 | 0.06428 | 0.04248 |
| 10 | 0.0696 | 0.040025 | 0.0401333 | 0.19244 | 0.17224 | 0.07694 | 0.03532 |
| 11 | 0.07054 | 0.038475 | 0.0459667 | 0.22674 | 0.20838 | 0.0934 | 0.04052 |
| 12 | 0.07242 | 0.039425 | 0.0452667 | 0.27084 | 0.25 | 0.1121 | 0.03434 |
| 13 | 0.07282 | 0.04055 | 0.0490333 | 0.31622 | 0.29468 | 0.13464 | 0.04178 |
| 14 | 0.07674 | 0.040975 | 0.0393333 | 0.3634 | 0.344 | 0.16006 | 0.0472 |
| 15 | 0.07486 | 0.040875 | 0.0415 | 0.40536 | 0.39502 | 0.19104 | 0.04482 |
| 16 | 0.07732 | 0.042425 | 0.0492333 | 0.4471 | 0.44884 | 0.24648 | 0.03988 |
| 17 | 0.08388 | 0.041825 | 0.0401 | 0.48894 | 0.47238 | 0.29144 | 0.04504 |
| 18 | 0.0835 | 0.042625 | 0.0410333 | 0.48846 | 0.48452 | 0.3039 | 0.04554 |
| 19 | 0.07834 | 0.041975 | 0.0408667 | 0.49728 | 0.5107 | 0.34228 | 0.03888 |
| 20 | 0.07888 | 0.041725 | 0.0429333 | 0.5007 | 0.53446 | 0.3837 | 0.03988 |
| 21 | 0.08162 | 0.0414 | 0.0462333 | 0.5055 | 0.52612 | 0.42072 | 0.04658 |
| 22 | 0.08138 | 0.038475 | 0.0425 | 0.51726 | 0.51706 | 0.44418 | 0.04026 |
| 23 | 0.08064 | 0.037325 | 0.0436333 | 0.5189 | 0.51166 | 0.46486 | 0.03544 |
| 24 | 0.0887 | 0.03745 | 0.0479 | 0.51486 | 0.50894 | 0.47512 | 0.02176 |
| 25 | 0.08918 | 0.03845 | 0.0454333 | 0.50946 | 0.50656 | 0.49934 | 0.02056 |
| 26 | 0.08404 | 0.03895 | 0.0474333 | 0.50364 | 0.50356 | 0.51024 | 0.02508 |
| 27 | 0.08756 | 0.0385 | 0.0502667 | 0.498 | 0.4985 | 0.5095 | 0.02954 |
| 28 | 0.0823 | 0.03885 | 0.0529667 | 0.49412 | 0.49294 | 0.50384 | 0.02034 |
| 29 | 0.08326 | 0.039575 | 0.0567667 | 0.49018 | 0.48692 | 0.4981 | 0.02272 |
| 30 | 0.08258 | 0.04055 | 0.0655 | 0.48516 | 0.48126 | 0.49344 | 0.02108 |
| 31 | 0.08294 | 0.04015 | 0.0809 | 0.4805 | 0.47426 | 0.48724 | 0.01858 |
| 32 | 0.0828 | 0.0403 | 0.0995667 | 0.47612 | 0.4665 | 0.47916 | 0.02664 |
| 33 | 0.08308 | 0.0412 | 0.1227 | 0.47324 | 0.46132 | 0.47074 | 0.01914 |
| 34 | 0.08638 | 0.04215 | 0.1511 | 0.46914 | 0.45658 | 0.4628 | 0.02018 |
| 35 | 0.0837 | 0.0414 | 0.189133 | 0.46566 | 0.44934 | 0.45758 | 0.02836 |
| 36 | 0.08398 | 0.0417 | 0.234733 | 0.46256 | 0.44678 | 0.45068 | 0.01458 |
| 37 | 0.08274 | 0.042775 | 0.273267 | 0.45844 | 0.44376 | 0.44406 | 0.02048 |
| 38 | 0.0829 | 0.043625 | 0.318867 | 0.45228 | 0.43978 | 0.43762 | 0.01374 |
| 39 | 0.08254 | 0.043025 | 0.361167 | 0.44952 | 0.43438 | 0.43214 | 0.0324 |
| 40 | 0.08756 | 0.043325 | 0.387667 | 0.44678 | 0.43082 | 0.42654 | 0.0157 |
| 41 | 0.0823 | 0.0441 | 0.4131 | 0.44378 | 0.42724 | 0.42194 | 0.01326 |
| 42 | 0.08222 | 0.044825 | 0.4464 | 0.44018 | 0.42334 | 0.41778 | 0.02214 |
| 43 | 0.08142 | 0.04365 | 0.477333 | 0.43758 | 0.41896 | 0.41392 | 0.01114 |
| 44 | 0.0833 | 0.0438 | 0.476867 | 0.43516 | 0.41546 | 0.4096 | 0.00884 |
| 45 | 0.08308 | 0.04475 | 0.4871 | 0.43238 | 0.4117 | 0.40626 | 0.0166 |
| 46 | 0.08522 | 0.045775 | 0.49 | 0.42896 | 0.40812 | 0.40358 | 0.02686 |
| 47 | 0.08 | 0.044225 | 0.494833 | 0.42526 | 0.40316 | 0.40116 | 0.01116 |
| 48 | 0.07956 | 0.04415 | 0.487833 | 0.42274 | 0.39962 | 0.39806 | 0.01202 |
| 49 | 0.07952 | 0.0453 | 0.4763 | 0.41952 | 0.39598 | 0.39578 | 0.00608 |
| 50 | 0.0788 | 0.0463 | 0.468967 | 0.41618 | 0.39234 | 0.39382 | 0.0104 |
| 51 | 0.0797 | 0.0449 | 0.4644 | 0.41282 | 0.38806 | 0.39208 | 0.01998 |
| 52 | 0.0795 | 0.0449 | 0.459667 | 0.40994 | 0.38454 | 0.38962 | 0.01684 |
| 53 | 0.07786 | 0.0459 | 0.4543 | 0.40704 | 0.38128 | 0.3883 | 0.0173 |
| 54 | 0.08232 | 0.0468 | 0.45 | 0.40346 | 0.3778 | 0.38674 | 0.00954 |
| 55 | 0.07642 | 0.045425 | 0.446933 | 0.40046 | 0.3742 | 0.385 | 0.00438 |
| 56 | 0.07612 | 0.04535 | 0.442567 | 0.39764 | 0.37128 | 0.38246 | 0.00326 |
| 57 | 0.07666 | 0.04645 | 0.436867 | 0.39508 | 0.36848 | 0.3809 | 0.0035 |
| 58 | 0.07414 | 0.046775 | 0.433 | 0.3913 | 0.3657 | 0.37906 | 0.00356 |
| 59 | 0.07408 | 0.045875 | 0.431433 | 0.38838 | 0.36254 | 0.37728 | 0.00296 |
| 60 | 0.07306 | 0.04575 | 0.427433 | 0.38592 | 0.36036 | 0.37436 | 0.00354 |
| 61 | 0.07262 | 0.046425 | 0.422767 | 0.38324 | 0.358 | 0.37222 | 0.00324 |
| 62 | 0.07138 | 0.04705 | 0.4209 | 0.37994 | 0.35606 | 0.37022 | 0.0033 |
| 63 | 0.07146 | 0.04575 | 0.418767 | 0.37752 | 0.35354 | 0.3685 | 0.00272 |
| 64 | 0.06956 | 0.045575 | 0.415933 | 0.3752 | 0.35188 | 0.3655 | 0.00438 |
| 65 | 0.06908 | 0.04685 | 0.411267 | 0.37322 | 0.35024 | 0.36426 | 0.00658 |
| 66 | 0.0684 | 0.047125 | 0.408367 | 0.37022 | 0.34864 | 0.3615 | 0.00732 |
| 67 | 0.06748 | 0.04605 | 0.4071 | 0.36806 | 0.34676 | 0.3585 | 0.0063 |
| 68 | 0.06668 | 0.046225 | 0.4036 | 0.36634 | 0.34552 | 0.3544 | 0.00376 |
| 69 | 0.06976 | 0.04645 | 0.3994 | 0.36428 | 0.34416 | 0.35084 | 0.00258 |

**Figure S7. Growth of MT123 under aerobic conditions without nitrate.** Cultures (n = 5 replicates) were grown at varying pH values. Points represent averaged data and error bars represent ±SD.

**Table S6. Growth statistics of *Castellaniella* isolates at a range of pHs**

| **Strain** | **pH** | **Growth Rate_MAX_ (hr^-1^)** | **Lag Time (h)** | **OD600_MAX_** |
| --- | --- | --- | --- | --- |
| MT123 | **5** | n.g. | n.g. | n.g. |
|  | **5.5** | 0.245* (0.015)** | 21.2 (2.4) | 0.533 (0.018) |
|  | **6** | 0.284 (0.026) | 13.5 (1.0) | 0.489 (0.023) |
|  | **7** | 0.360 (0.026) | 5.4 (0.9) | 0.456 (0.018) |
|  | **8** | 0.385 (0.026) | 5.0 (0.0) | 0.477 (0.007) |
| FW104-12G02 | **5** | 0.140 (0.096) | 28.7 (18.6) | 0.056 (0.052) |
|  | **5.5** | 0.640 (0.081) | 4.8 (1.3) | 0.135 (0.016) |
|  | **6** | 0.570 (0.047) | 3.0 (0.5) | 0.141 (0.004) |
|  | **7** | 0.513 (0.109) | 3.2 (0.0) | 0.096 (0.007) |
|  | **8** | 0.495 (0.110) | 3.8 (0.4) | 0.143 (0.027) |
| FW104-7C03 | **5** | 0.632 (0.121) | 12.6 (5.0) | 0.100 (0.017) |
|  | **5.5** | 0.741 (0.099) | 4.4 (0.5) | 0.158 (0.008) |
|  | **6** | 0.699 (0.097) | 4.6 (0.5) | 0.134 (0.008) |
|  | **7** | 0.599 (0.091) | 4.8 (0.5) | 0.099 (0.013) |
|  | **8** | 0.444 (0.082) | 4.2 (2.2) | 0.160 (0.040) |
| FW104-7G2B | **5** | 0.345 (0.198) | 20.8 (26.1) | 0.069 (0.057) |
|  | **5.5** | 0.614 (0.089) | 7.8 (0.5) | 0.229 (0.010) |
|  | **6** | 0.633 (0.041) | 6.0 (0.0) | 0.222 (0.007) |
|  | **7** | 0.671 (0.079) | 6.8 (1.5) | 0.066 (0.009) |
|  | **8** | 0.345 (0.048) | 4.1 (1.5) | 0.140 (0.027) |
| FW104-16D08 | **5** | n.g. | n.g. | n.g. |
|  | **5.5** | 0.296 (0.034) | 14.2 (1.4) | 0.039 (0.007) |
|  | **6** | 0.467 (0.049) | 10.2 (0.8) | 0.047 (0.009) |
|  | **7** | 0.415 (0.032) | 6.4 (0.5) | 0.058 (0.005) |
|  | **8** | 0.472 (0.072) | 7.5 (0.6) | 0.042 (0.009) |

*Average of n=5 replicates **±SD is shown in brackets

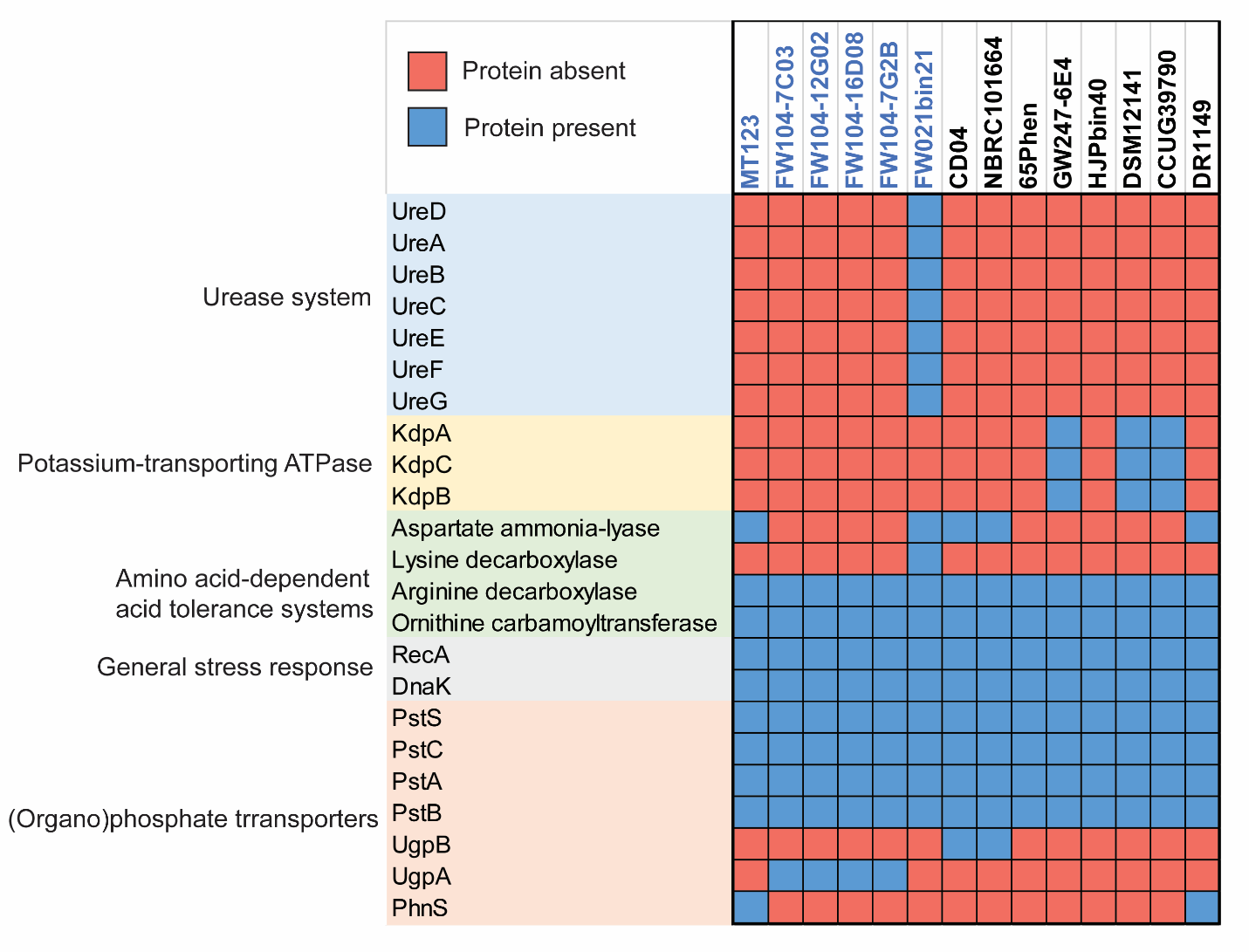

**Figure S8. Acid tolerance proteins encoded in *Castellaniella* genomes.** Presence (blue)/absence (red) matrix is shown for all *Castellaniella* genomes. ORR genomes are highlighted in blue. Proteins are grouped by their function in acid tolerance.

**Table S7. End-point nitrogen oxide measurements for MT123 and FW104-7G2B**

| **Strain/Condition** | **Initial OD** | **Final OD** | **Initial pH** | **Final pH** | **Final [NO_3_^-^] (mM)** | **Final [NO_2_^-^] (mM)** | **Final [N_2_O] (mM)** | **Estimated N_2_ (mM)** |
| --- | --- | --- | --- | --- | --- | --- | --- | --- |
| **MT123-pH 7** | 0.02* (0.00)** | 0.53 (0.01) | 6.59 (0.00) | 6.88 (0.02) | 0.05 (0.10) | 0.00 (0.00) | 0.00 (0.00) | 4.78 (0.05) |
| **MT123-pH 5.5** | 0.01 (0.00) | 0.48 (0.01) | 5.49 (0.00) | 6.06 (0.03) | 0.075 (0.05) | 0.00 (0.00) | 0.00 (0.00) | 4.80 (0.00) |
| **FW104-7G2B- pH 7** | 0.01 (0.00) | 0.27 (0.02) | 6.59 (0.00) | 6.91 (0.01) | 0.05 (0.06) | 0.00 (0.00) | 4.80 (0.07) | 0.00 (0.00) |

*Average of n=4 replicates **±SD is shown in brackets

**Table S8. Heavy metal homeostasis genes unique* to MT123**

| Heavy metal homeostasis genes | Metals |
| --- | --- |
| Cu(I)-responsive transcriptional regulator | Cu |
| Transcriptional regulator, ArsR family | Hg |
| Mg(2+) transport ATPase, P-type (EC 3.6.3.2) | Co Mg |
| 7-cyano-7-deazaguanine synthase (EC 6.3.4.20) | Al |
| Chromate reductase (EC 1.6.5.2) | Cr V |
| Arsenite/antimonite pump-driving ATPase ArsA (EC 3.6.3.16) | As Sb |
| Arsenic metallochaperone ArsD, transfers trivalent metalloids to ArsAB pump | As |
| FUPA30 P-type ATPase | Co Mg |
| Lead, cadmium, zinc, and mercury transporting ATPase (EC 3.6.3.3) (EC 3.6.3.5) | Cu Ag |
| Response regulator BaeR | Zn W |
| Sensory histidine kinase BaeS | Zn W |
| Mercuric transport protein, MerE | Hg |
| Mercuric resistance transcriptional repressor, MerD | Hg |
| Copper tolerance protein | Cu |
| Ferric iron ABC transporter, ATP-binding protein | Fe Ga |
| Mg(2+) transport ATPase, P-type (EC 3.6.3.2) | Co Mg |
| Magnesium and cobalt transport protein CorA | Mg Co Ni Mn |

*Defined as genes only observed in MT123 and not any of the other ORR *Castellaniella* isolates

OD600 (log-10)

| Time (h) | MT123 | MT123 COMM | FW104-12G02 | FW104-12G02 COMM | FW104-16D08 | FW104-16D08 COMM | FW104-7C03 | FW104-7C03 COMM | FW104-7G2B | FW104-7G2B COMM |
| --- | --- | --- | --- | --- | --- | --- | --- | --- | --- | --- |
| 0 | 0 | 0 | 0 | 0 |  | 0 | 0 | 0 | 0 | 0 |
| 1 | 0.0012 | 0.0024 | 0 | 0 |  | 0 | 0 | 0.0002 | 0 | 0.0034 |
| 2 | 0.0014 | 0.0044 | 0 | 0 |  | 0 | 0 | 0 | 0 | 0.0056 |
| 3 | 0.0044 | 0.0076 | 0.0052 | 0.0002 |  | 0 | 0 | 0 | 0.0006 | 0.0078 |
| 4 | 0.0088 | 0.0106 | 0.0176 | 0.0018 |  | 0 | 0.00075 | 0.001 | 0.0012 | 0.0108 |
| 5 | 0.013 | 0.0104 | 0.0298 | 0.0056 | 0 | 0 | 0.0035 | 0.003 | 0.0086 | 0.013 |
| 6 | 0.0202 | 0.017 | 0.0468 | 0.0082 | 0 | 0.0008 | 0.0095 | 0.0046 | 0.0176 | 0.0154 |
| 7 | 0.0226 | 0.0104 | 0.061 | 0.0084 | 0 | 0.0002 | 0.015 | 0.0044 | 0.0268 | 0.0146 |
| 8 | 0.0322 | 0.0136 | 0.08 | 0.0124 | 0 | 0.0004 | 0.02625 | 0.0078 | 0.0428 | 0.0152 |
| 9 | 0.0386 | 0.012 | 0.0938 | 0.0126 | 0 | 0.0006 | 0.042 | 0.0082 | 0.0584 | 0.0164 |
| 10 | 0.0466 | 0.0108 | 0.107 | 0.013 | 0.0004 | 0.0006 | 0.05575 | 0.0092 | 0.08 | 0.0174 |
| 11 | 0.0546 | 0.0088 | 0.1146 | 0.012 | 0.0016 | 0.0004 | 0.06925 | 0.009 | 0.0978 | 0.0158 |
| 12 | 0.0706 | 0.0118 | 0.124 | 0.0138 | 0.003 | 0.0012 | 0.08225 | 0.01 | 0.1172 | 0.0168 |
| 13 | 0.0856 | 0.0158 | 0.1296 | 0.0138 | 0.0076 | 0.0018 | 0.0915 | 0.012 | 0.136 | 0.019 |
| 14 | 0.1058 | 0.015 | 0.131 | 0.0126 | 0.0078 | 0.0012 | 0.095 | 0.0106 | 0.1502 | 0.0188 |
| 15 | 0.1352 | 0.0208 | 0.1358 | 0.0136 | 0.0116 | 0.0016 | 0.0985 | 0.0132 | 0.1654 | 0.0194 |
| 16 | 0.17 | 0.0242 | 0.1366 | 0.0116 | 0.0138 | 0.0008 | 0.1 | 0.012 | 0.1768 | 0.0192 |
| 17 | 0.213 | 0.0282 | 0.139 | 0.0136 | 0.0174 | 0.001 | 0.10275 | 0.0128 | 0.1892 | 0.0204 |
| 18 | 0.2568 | 0.0374 | 0.1428 | 0.0146 | 0.0196 | 0.0024 | 0.10525 | 0.0148 | 0.1978 | 0.0216 |
| 19 | 0.304 | 0.0446 | 0.1418 | 0.0146 | 0.0232 | 0.0026 | 0.10525 | 0.0164 | 0.2058 | 0.0204 |
| 20 | 0.3674 | 0.055 | 0.1418 | 0.0134 | 0.0238 | 0.002 | 0.10575 | 0.0148 | 0.2112 | 0.0198 |
| 21 | 0.4036 | 0.0726 | 0.1454 | 0.0168 | 0.029 | 0.004 | 0.109 | 0.0188 | 0.219 | 0.024 |
| 22 | 0.438 | 0.0898 | 0.1454 | 0.014 | 0.028 | 0.005 | 0.108 | 0.019 | 0.2206 | 0.0226 |
| 23 | 0.4348 | 0.113 | 0.1468 | 0.0144 | 0.0294 | 0.006 | 0.109 | 0.0218 | 0.2224 | 0.0222 |
| 24 | 0.4346 | 0.1432 | 0.1494 | 0.0174 | 0.032 | 0.0078 | 0.11225 | 0.024 | 0.2254 | 0.0248 |
| 25 | 0.4306 | 0.1662 | 0.149 | 0.0136 | 0.0302 | 0.0056 | 0.10825 | 0.0208 | 0.223 | 0.0218 |
| 26 | 0.432 | 0.2088 | 0.151 | 0.0166 | 0.0312 | 0.0074 | 0.11175 | 0.0242 | 0.2246 | 0.023 |
| 27 | 0.4308 | 0.26 | 0.151 | 0.0172 | 0.0316 | 0.0068 | 0.11075 | 0.0262 | 0.2228 | 0.0248 |
| 28 | 0.4304 | 0.307 | 0.1508 | 0.0172 | 0.034 | 0.0078 | 0.11 | 0.027 | 0.225 | 0.0242 |
| 29 | 0.4294 | 0.3446 | 0.152 | 0.0166 | 0.031 | 0.008 | 0.11175 | 0.0248 | 0.2254 | 0.0232 |
| 30 | 0.4334 | 0.3704 | 0.1532 | 0.0196 | 0.0336 | 0.009 | 0.11275 | 0.03 | 0.229 | 0.026 |
| 31 | 0.4314 | 0.389 | 0.1516 | 0.0174 | 0.033 | 0.009 | 0.113 | 0.0286 | 0.2276 | 0.0236 |
| 32 | 0.4298 | 0.3962 | 0.1516 | 0.0184 | 0.0312 | 0.009 | 0.11275 | 0.0314 | 0.227 | 0.0252 |
| 33 | 0.4322 | 0.4078 | 0.151 | 0.0188 | 0.032 | 0.0106 | 0.11475 | 0.0326 | 0.2278 | 0.0258 |
| 34 | 0.43 | 0.4106 | 0.1528 | 0.0204 | 0.0348 | 0.0094 | 0.11375 | 0.0316 | 0.2278 | 0.0264 |
| 35 | 0.429 | 0.4138 | 0.1502 | 0.0188 | 0.0336 | 0.0086 | 0.11375 | 0.0346 | 0.2284 | 0.0238 |
| 36 | 0.4282 | 0.4208 | 0.1514 | 0.0202 | 0.0324 | 0.0102 | 0.114 | 0.0362 | 0.2268 | 0.0244 |
| 37 | 0.4282 | 0.4238 | 0.1502 | 0.0216 | 0.0338 | 0.0102 | 0.1135 | 0.0378 | 0.228 | 0.0262 |
| 38 | 0.427 | 0.4224 | 0.1514 | 0.0218 | 0.0338 | 0.0124 | 0.11375 | 0.0402 | 0.2284 | 0.0282 |
| 39 | 0.4298 | 0.431 | 0.1524 | 0.0222 | 0.0342 | 0.0124 | 0.11675 | 0.0408 | 0.2288 | 0.0286 |
| 40 | 0.4292 | 0.435 | 0.1516 | 0.0236 | 0.034 | 0.013 | 0.11575 | 0.0398 | 0.2292 | 0.0288 |
| 41 | 0.4296 | 0.4336 | 0.1504 | 0.0236 | 0.0322 | 0.0116 | 0.11475 | 0.0386 | 0.2286 | 0.0282 |
| 42 | 0.4304 | 0.4376 | 0.1504 | 0.024 | 0.0336 | 0.0122 | 0.11475 | 0.0404 | 0.2298 | 0.0284 |
| 43 | 0.4308 | 0.437 | 0.154 | 0.0258 | 0.0328 | 0.0122 | 0.11475 | 0.0404 | 0.2284 | 0.029 |
| 44 | 0.4328 | 0.4442 | 0.1532 | 0.0274 | 0.0348 | 0.0138 | 0.11825 | 0.0452 | 0.231 | 0.0308 |
| 45 | 0.4314 | 0.4472 | 0.1518 | 0.0264 | 0.0338 | 0.0144 | 0.11775 | 0.0438 | 0.2292 | 0.0322 |
| 46 | 0.4306 | 0.4438 | 0.1502 | 0.0286 | 0.0362 | 0.0142 | 0.11675 | 0.046 | 0.2292 | 0.0306 |
| 47 | 0.4306 | 0.4464 | 0.1506 | 0.0276 | 0.0354 | 0.0138 | 0.11575 | 0.0448 | 0.2296 | 0.0296 |
| 48 | 0.4326 | 0.452 | 0.1522 | 0.0312 | 0.037 | 0.0146 | 0.118 | 0.047 | 0.2302 | 0.0322 |
| 49 | 0.4302 | 0.4502 | 0.15 | 0.0304 | 0.0368 | 0.015 | 0.117 | 0.0486 | 0.2292 | 0.0318 |
| 50 | 0.43 | 0.449 | 0.1508 | 0.0334 | 0.0364 | 0.0148 | 0.11675 | 0.0494 | 0.229 | 0.0324 |
| 51 | 0.4306 | 0.4518 | 0.1506 | 0.0336 | 0.0372 | 0.015 | 0.11975 | 0.0488 | 0.231 | 0.0332 |
| 52 | 0.4314 | 0.4552 | 0.1524 | 0.0368 | 0.0388 | 0.0166 | 0.12075 | 0.051 | 0.2326 | 0.0326 |
| 53 | 0.431 | 0.458 | 0.1518 | 0.0376 | 0.0384 | 0.0166 | 0.11975 | 0.0524 | 0.2314 | 0.0334 |
| 54 | 0.4352 | 0.4606 | 0.1526 | 0.04 | 0.0412 | 0.0174 | 0.12 | 0.0542 | 0.2314 | 0.0352 |
| 55 | 0.4344 | 0.4594 | 0.1504 | 0.0386 | 0.0396 | 0.0144 | 0.11875 | 0.0522 | 0.2294 | 0.0328 |
| 56 | 0.435 | 0.4646 | 0.1484 | 0.04 | 0.0392 | 0.016 | 0.1185 | 0.0518 | 0.2308 | 0.0338 |
| 57 | 0.4352 | 0.4636 | 0.1488 | 0.0386 | 0.0392 | 0.0166 | 0.119 | 0.053 | 0.2308 | 0.032 |
| 58 | 0.438 | 0.466 | 0.1488 | 0.039 | 0.0392 | 0.016 | 0.1205 | 0.0554 | 0.2316 | 0.0342 |
| 59 | 0.4406 | 0.4654 | 0.1508 | 0.0428 | 0.0402 | 0.018 | 0.12125 | 0.0566 | 0.2316 | 0.0376 |
| 60 | 0.4436 | 0.4754 | 0.1512 | 0.0448 | 0.0442 | 0.0196 | 0.1225 | 0.0594 | 0.237 | 0.0384 |
| 61 | 0.4438 | 0.4696 | 0.1498 | 0.0444 | 0.0424 | 0.0174 | 0.12325 | 0.058 | 0.232 | 0.0376 |
| 62 | 0.4414 | 0.4772 | 0.1468 | 0.0412 | 0.0408 | 0.0152 | 0.11925 | 0.0566 | 0.2306 | 0.035 |
| 63 | 0.4464 | 0.4768 | 0.1496 | 0.046 | 0.0428 | 0.019 | 0.122 | 0.0614 | 0.2344 | 0.0398 |
| 64 | 0.4496 | 0.4832 | 0.1494 | 0.0466 | 0.0436 | 0.0182 | 0.12275 | 0.0612 | 0.2364 | 0.039 |
| 65 | 0.4492 | 0.4824 | 0.1498 | 0.049 | 0.0442 | 0.021 | 0.123 | 0.062 | 0.2354 | 0.04 |
| 66 | 0.4496 | 0.4846 | 0.148 | 0.0474 | 0.0436 | 0.0188 | 0.12175 | 0.0598 | 0.236 | 0.0378 |
| 67 | 0.4496 | 0.4852 | 0.1488 | 0.0506 | 0.0442 | 0.02 | 0.12325 | 0.0648 | 0.2356 | 0.0392 |
| 68 | 0.45 | 0.4866 | 0.1478 | 0.0502 | 0.0424 | 0.017 | 0.12175 | 0.0654 | 0.2362 | 0.038 |
| 69 | 0.4468 | 0.491 | 0.1484 | 0.0526 | 0.0444 | 0.0196 | 0.12375 | 0.0668 | 0.239 | 0.04 |
| 70 | 0.4534 | 0.4914 | 0.149 | 0.0524 | 0.0442 | 0.0186 | 0.126 | 0.07 | 0.2348 | 0.0408 |
| 71 | 0.4526 | 0.4964 | 0.1458 | 0.051 | 0.0432 | 0.0172 | 0.12175 | 0.0674 | 0.237 | 0.0376 |
| 72 | 0.453 | 0.503 | 0.1472 | 0.0532 | 0.0436 | 0.019 | 0.12525 | 0.0706 | 0.2374 | 0.0388 |

**Figure S9. Growth of the ASV1 and ASV2 representative strains +/- FW104 COMM.** The composition of the FW104 COMM represents the FW104 groundwater. pH was adjusted to 5.5 to reflect the average pH of the FW104 groundwater. Filled circles represent control cultures without the added metal mixture. Unfilled circles represent FW104 COMM-exposed cultures. Individual points reflect the average of five replicates. Error bars represent ±SD.
